# Sex-biased zinc responses modulate ribosome biogenesis, protein synthesis and social defects in *Cttnbp2* mutant mice

**DOI:** 10.1101/2025.02.03.636382

**Authors:** Yu-Lun Fang, Tzu-Li Yen, Hou-Chen Liu, Ting-Fang Wang, Yi-Ping Hsueh

## Abstract

Autism spectrum disorders (ASD) are neurodevelopmental conditions influenced by genetic mutations, dietary factors, and sex-specific mechanisms, yet the interplay of these factors remains elusive. Here, we investigate the sex-biased responses of mutant mice carrying an ASD-associated mutation in *Cttnbp2* to dietary zinc supplementation using behavioral assays, proteomic and bioinformatic analyses, and puromycin pulse labeling to assess protein synthesis. Our results demonstrate that zinc supplementation enhances ribosome biogenesis and increases the density and size of dendritic spines in male *Cttnbp2* mutant mice, alleviating male-biased social deficits. Analyses of neuronal cultures further revealed that neurons, not astrocytes, respond to zinc to enhance protein synthesis. In contrast, female *Cttnbp2* mutants exhibit resilience to differential zinc intake, even under zinc deprivation. Elevated mTOR phosphorylation and increased protein levels of translational initiation factors in female brains may provide a protective mechanism, reducing their sensitivity to zinc deficiency. *Cttnbp2* mutations heighten male vulnerability to zinc deprivation, impairing social behaviors. These findings highlight zinc-regulated ribosome biogenesis and protein synthesis as critical mediators of sex-specific ASD phenotypes, offering new insights into dietary interventions.

## Introduction

Autism spectrum disorders (ASD) are a group of common neurodevelopmental conditions. As of 2020, approximately 1 in 36 children aged 8 years in the USA has been diagnosed with ASD (source: <u>CDC</u>). ASD exhibits a pronounced male bias, with a boy-to-girl ratio of about 4:1, meaning roughly 4% of boys and 1% of girls are affected (Maenner *et al*., 2023). The condition has been linked to hundreds of genes (SFARI gene database, https://gene.sfari.org/database/human-gene/). Additionally, environmental factors interact with genetic variations to influence ASD outcomes (Lee *et al*., 2022b; Lin *et al*., 2022; Lu and Hsueh, 2022; Mohebalizadeh *et al*., 2023; Mony and Paoletti, 2023), further complicating research into its etiology.

Zinc, an essential nutrient, is highly relevant to ASD (Hsueh, 2025). ASD patients tend to have lower zinc levels than normal based on analyses of their blood, hair, and teeth (Alsufiani *et al*., 2022; Arora *et al*., 2017; Pfaender *et al*., 2017; Yasuda *et al*., 2011). Notably, men require 30-50% more zinc than women to maintain optimal health (https://ods.od.nih.gov/HealthInformation/ nutrientrecommendations.aspx). As a structural or enzymatic cofactor, zinc is critical for numerous proteins and plays essential roles in cell growth, differentiation and metabolism across various cell types (Brown *et al*., 2002). Among zinc-binding proteins, zinc finger proteins are the most abundant and well-studied, encoded by up to 3% of genes in the human genome (Klug, 2010). These proteins regulate a wide range of molecular functions, including transcription, DNA repair, ubiquitin-dependent protein degradation, and signal transduction (Cassandri *et al*., 2017; Klug and Rhodes, 1987).

In addition to its ubiquitous functions in all cell types, zinc is especially critical in the brain. The average concentration of zinc in mammalian brains is 150 μM, which is 10-fold higher than the level circulating in serum (Mocchegiani *et al*., 2005; Portbury and Adlard, 2017; Takeda, 2001). Approximately 10-15% of the zinc in brains is concentrated in synaptic vesicles and it is co-released with glutamate upon synaptic stimulation, playing an essential role in modulating neurotransmission (Cole *et al*., 1999; Frederickson, 1989; Frederickson *et al*., 2000; Huang, 1997; Lee *et al*., 2015; Tóth, 2011). Recent studies have further indicated that zinc regulates the protein-protein interactions and condensate formation of postsynaptic proteins, and also controls synaptic targeting and retention of these proteins (Baron *et al*., 2006; Grabrucker *et al*., 2014; Shih *et al*., 2022; Shih *et al*., 2020a).

Several ASD-causative genes have been shown to bind zinc or to be regulated by it (Grabrucker, 2014; Grabrucker *et al*., 2014; Shih *et al*., 2022; Shih *et al*., 2020a; Shih *et al*., 2020b; Vyas *et al*., 2021; Vyas *et al*., 2020). Among them, cortactin-binding protein 2 (CTTNBP2), a zinc-binding cytoskeleton-associated protein (Shih *et al*., 2022), is specifically expressed in neurons (Chen and Hsueh, 2012; Hu *et al*., 2016; Ohoka and Takai, 1998). Due to alternative splicing, multiple transcript variants of the *Cttnbp2* are predicted (Chen and Hsueh, 2012). However, only the short *Cttnbp2* transcript, encoding a 630-amino-acid protein, is expressed in the brain (Chen and Hsueh, 2012; Shih *et al*., 2020a). Hereafter, we refer to this short isoform as CTTNBP2. Given its dominant expression in neurons, mutations in the short form of *Cttnbp2* are particularly relevant to ASD and neuronal function, especially synapse formation (Shih *et al*., 2020b).

CTTNBP2 is highly enriched at dendritic spines via condensate formation of liquid-liquid phase separation and is critical for regulating dendritic spine formation and maintenance (Chen and Hsueh, 2012; Shih *et al*., 2022; Shih *et al*., 2020a; Shih *et al*., 2020b). CTTNBP2 interacts with zinc through its N-terminal coiled-coil region (Shih *et al*., 2022). Zinc binding induces CTTNBP2 to undergo a liquid-to-gel phase transition, immobilizing the protein and enhancing its synaptic retention (Shih *et al*., 2022). In addition to being influenced by zinc, CTTNBP2 also regulates zinc levels in the brain. *Cttnbp2* knockout in mice reduces total zinc levels in their brains and impairs the synaptic distribution of several zinc-related and autism-associated proteins (Shih *et al*., 2020a). An increase in dietary zinc intake for a week was previously found to restore zinc levels in the mutant mice’s brain and enhance the synaptic distribution of CTTNBP2-regulated proteins (Shih *et al*., 2020a). Thus, CTTNBP2 binds zinc and contributes to zinc homeostasis in the brain.

Consistent with genetic analyses of patients with ASD (De Rubeis *et al*., 2014), we previously shown that knockout and ASD-linked mutations of the *Cttnbp2* gene elicit ASD-related behaviors in mice, including reduced social interaction, cognitive inflexibility, and impaired ultrasonic vocalization (Shih *et al*., 2020a). Similarly, paralleling findings that CTTNBP2 mutations are observed only in male, but not female, patients with ASD (De Rubeis *et al*., 2014; Sanders *et al*., 2015), we showed that *Cttnbp2* deficiency—through knockout or ASD-linked mutations—results in social deficits in male, but not female, mice when fed a 30 or 84 ppm zinc diet (Yen *et al*., 2023). Importantly, feeding male *Cttnbp2^–/–^*, *Cttnbp2^+/M120I^*and *Cttnbp2^+/R533*^* mice a 150 ppm zinc diet for seven days improved their social behaviors (Shih *et al*., 2020a; Shih *et al*., 2020b), further supporting the beneficial effect of zinc supplementation on brain function.

Continuous zinc supplementation is required to maintain the beneficial effects of zinc to male *Cttnbp2*-deficient mice, as our previous studies showed that the effect of zinc supplementation diminished within seven days after discontinuing additional zinc intake (Shih *et al*., 2022; Shih *et al*., 2020a). However, the impact of zinc deprivation and constant zinc supplementation on *Cttnbp2*-deficient mice remains unclear. Here, we dissect the sex-differential responses of *Cttnbp2* mutant mice to long-term changes in dietary zinc. We chose to focus on *Cttnbp2^+/M120I^* mice as a model because the M120I mutation disrupts several important molecular functions of CTTNBP2, including N- and C-terminal interactions, binding of cortactin and liquid-liquid phase separation (Shih *et al*., 2022; Shih *et al*., 2020b), consequently impairing the dendritic spine retention of CTTNBP2 and dendritic spine formation by neurons (Shih *et al*., 2022; Shih *et al*., 2020b; Yen *et al*., 2023). However, the CTTNBP2 M120I mutant protein can still bind zinc (Shih *et al*., 2022), and notably, the phenotypes of *Cttnbp2^+/M120I^* mice were improved by zinc supplementation for 7 days (Shih *et al*., 2022; Yen *et al*., 2023). In this report, we designed three sets of experiments—including behavioral assays, synaptic proteome analysis, and evaluation of protein synthesis—to assess the sex-differential responses of *Cttnbp2^+/M120I^*mice to zinc supplementation. Our results indicate that in addition to acting on synapses, long-term dietary zinc supplementation differentially controls ribosome biogenesis and protein synthesis, likely via transcriptional regulation, resulting in the observed sex bias in *Cttnbp2*-related disorders.

## Materials and Methods

### Experimental model and subject details

The *Cttnbp2^M120I/+^* mice have been generated and characterized previously (Shih *et al*., 2020a; Shih *et al*., 2020b; Yen *et al*., 2023). The mouse line was maintained by backcrossing to C57BL/6JNarl mice purchased from the National Laboratory Animal Center, Taiwan. Mice were housed in the animal facility of the Institute of Molecular Biology, Academia Sinica, under controlled temperature (20-23 °C) and humidity (48%–55%) and a 12 h light/12 h dark cycle (light-dark: 8:00 am-8:00 pm) with free access to water and food. To maintain the mouse line, LabDiet 5K54 containing 84 ppm of zinc and 19% protein was provided. For special zinc diet experiments, the following diets were considered: (1) 30 ppm zinc, i.e., Research Diets D19410B, representing a normal zinc diet; (2) 150 ppm zinc: Research Diets D19410B plus 75 ppm ZnSO_4_ in drinking water, representing a higher zinc diet; (3) 30-ppm deprivation: mice first fed on 30 ppm zinc diet and then switched to Research Diets D19401 (0.85 ppm zinc) from P28; (4) 150-ppm deprivation: mice first fed on 150 ppm zinc diet and then switched to the 0.85 ppm zinc diet from P28. Behavioral assays and body weight measurements were performed during P28-P42. For behavior assays, both male and female mice were moved from the breeding room to the experimental area and housed individually after weaning at P21-22. Temperature and humidity in the test room were also controlled and the light intensity was 240 lumen/m^2^ (lux). All experiments were performed during the daytime, from 10.00 am ∼ 5.00 pm. All animal experiments were performed with the approval of the Academia Sinica Institutional Animal Care and Utilization Committee (Protocol No. 11-12-294 and 23-03-1990) and in strict accordance with its guidelines, those of the Council of Agriculture Guidebook for the Care and Use of Laboratory Animals, and ARRIVE.

### Experimental design

We used *Cttnbp2^+/M120I^* mice (Shih *et al*., 2020b; Yen *et al*., 2023) as an ASD mouse model to investigate the effect of long-term dietary zinc supplementation on dendritic spine formation, synaptic proteomes and social behaviors. Both male and female mice were used to study the sex-biased response of *Cttnbp2^+/M120I^* mice. The underlying molecular mechanism was explored using bioinformatic analyses, immunoblotting, and puromycin labeling for protein synthesis.

### Antibodies and reagents

The following antibodies were used in this study: GFP (Abcam, ab13970, chicken, 1:1000); RPL10 (Novus Biologicals, NBP1-84037, rabbit, 1:5000); RPS29 (Proteintech, 17374-1-AP, rabbit, 1:2000); COX6A1 (Proteintech, 11460-1-AP, rabbit, 1:1000); TOMM22 (Invitrogen, PA5-89898, rabbit, 1:3000); DARPP32 (Chemicon, AB1656, rabbit, 1:1000); HSP90 serum (gift from Dr. Chung Wang, rabbit, 1:2000) (Liou and Wang, 2005); p-mTOR (Cell Signaling, 5536S, rabbit, 1:1000); mTOR (Cell Signaling, 2972S, rabbit, 1:500); puromycin (Millipore, MABE343, 1:2000); GAPDH (ProteinTech, 10494-1-AP, 1:5000); EIF1 (Cell Signaling, 12496S, 1:1000); EIF1a (Abcam, EPR12466(B), 1:5000); horseradish peroxidase-conjugated anti-mouse IgG antibody (ProteinTech, SA00001-1, 1:5000); horseradish peroxidase-conjugated anti-rabbit IgG antibody (ProteinTech, SA00001-2, 1:5000); Alexa Fluor-488-conjugated anti-chicken IgY antibody (Invitrogen, A-11039, 1:1000). The following reagents were obtained commercially: OCT (Tissue-TeK, 4583); DAPI (Invitrogen, D1306, 1:1000); ZnSO_4_ (Sigma-Aldrich, #Z0251); NH_4_HCO_3_ (09830, Fluka); methanol (1.06009.2500, Merck); acetonitrile (51101, Pierce); DTT (43815, Fluka); sequencing grade modified trypsin (V5111, Promega); formic acid (1.00264.1000, Merck); Zip-Tips (Millipore, ZTC 18M 096); 30 ppm zinc diet (Research Diets, D19410B); 0.85 ppm zinc diet (Research Diets, D19401); Bradford assay (Bio-Rad Protein Assay Dye Reagent Concentrate, Cat #5000006); WesternBright ECL Spray (Advansta Inc., K-12049-D50); Immobilon Western Chemiluminescent HRP Substrate (Millipore, WBKLS0050); Brilliant Blue R (Sigma-Aldrich, B7920); 50× MEM amino acids solution (Gibco, 11130-051); 100× MEM non-essential amino acids solution (Gibco, 11140-050); 1× DMEM (Gibco, 11960-044); Neurobasol medium (Gibco, 21103-049); Pen/Strep (Gibco, 15140-122); 100× L-glutamine 200 mM (Gibco, A29168-01); 50× B-27 supplement (Gibco, 17504-044); and Papain from papaya latex lyophilized powder (Sigma-Aldrich, P4762-500MG).

### The association of SFARI genes with zinc and sex

All 1191 SFARI genes (https://gene.sfari.org/database/human-gene/) from the August 19, 2024 release were analyzed based on annotations in UniProt (https://www.uniprot.org) from September 5, 2024. The genes containing “zinc” in the sections of “Function”, “Keyword” and “Gene ontology” of the UniProt database were selected as zinc-related SFARI genes (**Supplementary Table S1**). To search for sex-related genes, SFARI genes containing the keywords “estrogen”, “androgen”, “sex”, “male”, “female” and “thyroxine” in the sections of “Function”, “Keyword” and “Gene ontology” of the UniProt database were selected (**Supplementary Tables S2, S3**).

### Immunostaining

Fifty-μm-thick brain sections of the ILA region were prepared for immunofluorescence staining as described previously (Lin *et al*., 2023; Yen *et al*., 2023). After serial washing using regular and high-salt TBS (25 mM Tris pH 7.5, regular: 0.85% NaCl; high-salt: 1.7% NaCl), brain sections were permeabilized with 0.06% Triton X-100 in high-salt TBS (i.e., TBST) for 10 min and blocked with blocking buffer (2% BSA and 3% horse serum in regular TBST) for 1 hr. Chicken anti-GFP primary antibody was added into the blocking buffer and incubated with brain sections overnight at 4 °C. After washing with high-salt TBST, brain sections were then incubated with Alexa Fluor-488-conjugated anti-chicken IgY secondary antibody and DAPI in the blocking buffer for 2 hr. After washing, signals were acquired using a confocal LSM 700 microscope (Carl Zeiss) with an immersion objective (iPlan-Apochromat 63X/1.40 Oil DIC M27) with Zen 2009 version 6.0 software, and then manually quantitated in ImageJ (Lin *et al*., 2023). To quantify the morphology of dendritic spines in the ILA, the primary branches of apical dendrites with the branch point located ∼100-250 μm from the soma were selected. The dendritic segments of 20 μm in length starting from a point 5 µm away from the branch point were selected for quantification.

### Sample preparation for liquid chromatography-mass spectrometry (LC-MS-MS)

The brain samples were prepared as described previously (Shih *et al*., 2020a; Shih *et al*., 2020c; Yen *et al*., 2023). In brief, one hemisphere of the cortex from 6-week-old mice was isolated and homogenized using a tissue Dounce homogenizer with a loose pestle in 1 ml buffered sucrose (10 mM HEPES pH 7.5, 320 mM sucrose, 10 mM DTT, 5 mM EDTA, 1 mM EGTA, 10 mM β-glycerophosphate, 2 mM MgCl_2_, 2 mg/ml leupeptin, 2 mg/ml pepstatin-A, 2 mg/ml aproteinin, 2 mM PMSF, 2 mg/ml MG132). Total homogenate was centrifuged at 800 × g for 10 min at 4 °C. The supernatant was collected and centrifuged again under the same condition to ensure the supernatant was well separated from the pellets. The supernatant was then centrifuged at 9200 × g for 15 min to collect the pellet as a crude synaptosomal fraction. The protein concentrations of crude synaptosomal fractions were determined by Bradford assay. Synaptosomal protein samples were then analyzed with LC-MS-MS and immunoblotting.

### LC-MS-MS

LC-MS-MS was conducted as described previously (Shih *et al*., 2020a; Shih *et al*., 2020c; Yen *et al*., 2023). In brief, synaptosomal protein samples were prepared from three mice for each group. Five μg of protein was loaded for 4% SDS-PAGE. After the dye front had completely entered the gel (∼10 min), the gel was fixed with 50% methanol and 10% acetic acid for 10 min, and stained for a further 10 min with 0.1% Brilliant Blue R (0.1% Brilliant Blue R, 50% methanol, 10% acetic acid) followed by destaining with 10% methanol and 7% acetic acid until the bands were clearly visible. The gel bands were excised and cut into small pieces for a modified in-gel digestion protocol using trypsin (Shevchenko *et al*., 2006). Extracted peptide products were desalted using Zip-Tips before mass spectrometric analysis. The nanoLC-nanoESI-MS/MS analysis was performed using an EASY-nLC 1200 system connected to a Thermo Orbitrap Fusion Lumos mass spectrometer (Thermo-Fisher Scientific, Bremen, Germany) equipped with a Nanospray Flex ion source (Thermo-Fisher Scientific, Bremen, Germany) as described previously (Shih *et al*., 2020a).

### Proteomic analysis and label-free quantification

The bioinformatics analysis of LC-MS-MS data was performed as described previously (Shih *et al*., 2020a; Shih *et al*., 2020c; Yen *et al*., 2023). Our proteomic data were searched against the Swiss-Prot *Mus musculus* database (17,049 entries total) using the Mascot search engine (v.2.6.2; Matrix Science, Boston, MA, USA) through Proteome Discoverer (v 2.2.0.388; Thermo-Fisher Scientific, Waltham, MA, USA). The search criteria were trypsin digestion, carbamidomethyl (C) for the fixed modification, oxidation (M) and acetylation (protein N-terminal) for the variable modifications, two missed cleavages, and 10 ppm of mass accuracy for the parent ion and 0.6 Da for the fragment ions. The false discovery rate (FDR) was calculated using the Proteome Discoverer Percolator function, and identifications with an FDR > 1% were rejected. For label-free quantification, precursor ion areas were extracted using the Minora Feature Detector node in Proteome Discoverer 2.2 with a 2 ppm mass precision and 2 min retention time-shift (to align the LC/MS-MS peaks mapping the isotope pattern and retention time). The ratio for each peptide was normalized against the total identified peptides and was used for protein-level quantification.

### Analyses of functional protein networks and gene ontology

Differentially expressed proteins (DEPs) were selected based on two criteria: an adjusted p-value < 0.05 and reliable peptide quality. Gene ontology (GO) of DEPs was conducted using the enrichment bubble plot feature of SRplot (https://www.bioinformatics.com.cn/plot_basic_gopathway_enrichment_bubbleplot_081_en).

STRING analysis (version 11.0, https://string-db.org/) was employed to identify GO terms and functional protein networks. Lines between nodes in our figures indicate that the interactions are based on experimental or STRING database evidence (https://string-db.org). The thickness of the lines indicates the confidence associated with the interaction. Different colors of nodes indicate associations with various gene ontology (GO), reactome (MMU) or annotated keyword (KW) pathways. Some nodes have multiple colors, indicating their involvement in multiple pathways. SynGO analysis (https://www.syngoportal.org/) was used to analyze protein enrichment at synapses.

### Primary cortical neuronal culture and zinc treatment

The procedures for primary neuronal culture have been described previously (Hung *et al*., 2018). In brief, mixed-sex embryonic mouse cortex was collected at embryonic day 18 (E18) and digested with papain solution (0.6 mg/ml papain, 0.5 mM EDTA, 1.5 mM CaCl_2_, 0.06% DNaseI, 0.2 mg/ml cysteine) at 37 °C for 15 min. The papain solution was then washed out three times using pre-warmed neural maintenance medium [50% Neurobasal medium, 50% DMEM, 1% Pen-Strep (Gibco), 2 mM Glutamine (Gibco)], followed by gentle pipetting to dissociate cortical neurons in 10 ml of neural maintenance medium and centrifugation at 900 rpm for 5 min. The cell pellets were resuspended in fresh pre-warmed neural maintenance medium with 1× essential and non-essential amino acids and seeded on poly-L-lysine (1 mg/ml)-precoated 12 well culture plates.

#### Long-term zinc treatment

One hour after plating, cells attached to culture plate or coverslips. An additional 100 µM ZnSO_4_ was added to the medium until the neurons were subjected to further experiments.

#### Short-term zinc treatment

An additional 100 µM ZnSO_4_ was added to the medium at 16 DIV. Two days later, neurons were subjected to further experiments.

### Puromycin treatment and neuronal harvesting

Puromycin pulse labeling was performed as described previously (Shih and Hsueh, 2016). At 10 or 18 DIV, 10 µg/ml puromycin was added into the neural culture medium for pulse labeling for 10 min. After removing the puromycin-containing medium, the conditioned medium containing ¼ fresh medium was added back to cultured neurons and incubated for an additional 50 min. At the end of incubation, neurons were washed twice with ice-cold PBS and collected in SDS sample buffer using a cell scraper. The cell lysates were then boiled for 10 min followed by a 10-sec vortex.

### Immunoblotting

#### Brain lysates

Immunoblotting using brain lysates was conducted as described previously (Shih *et al*., 2020a). Three biological replicates (10 or 15 µg per lane) of cortical synaptosomal lysates were subjected to independent immunoblotting assays to validate the results of our proteomic analysis and to evaluate the effect of zinc supplementation on *Cttnbp2^+/M120I^* mice. The membranes were blocked with 10% non-fat milk, washed with PBST (PBS with 0.2% Tween-20) buffer, and then incubated with primary antibodies overnight at 4 °C. After washing, the membrane was incubated with HRP-conjugated secondary antibodies for 1 h at room temperature. HSP90 and GAPDH was used as an internal control to normalize the protein levels. The results were visualized using a WesternBright ECL Spray or Immobilon Western Chemiluminescent HRP Substrate and recorded using an ImageQuant LAS 4000 system with ImageQuant LAS 4000 Biomolecular Imager software (GE Healthcare). Quantification was performed using ImageJ Fiji version 2.1.0/1.53t.

#### Neuronal culture lysates

To evaluate the effect of zinc on protein synthesis and ribosome biogenesis, 100 μl SDS sample buffer was added into each well of the 12-well culture plate to lyse cells. Equal fractions (10 µl per lane) of three experimental replicates of neuronal lysate from one batch (from a total of five independent batches) were loaded into SDS-PAGE for Coomassie blue staining and immunoblotting assay. To assess total protein amounts after zinc treatment, the SDS-PAGE gels were stained with Coomassie blue solution (0.1% Brilliant Blue R) for 30 min, then destained by destaining buffer until the bands were clearly visible. The results were recorded by means of digitalization of trans-illumination using an ImageQuant LAS 4000 system. Immunoblotting assay and quantification protocols for puromycin and ribosome proteins are described above.

### Animal behavior assays

In addition to measuring body weight, four behavioral tests, i.e., open field test (OF), elevated plus maze (EPM), three-chamber test (3C) and reciprocal social interaction (RSI), were conducted sequentially starting at P28 and ending at P42, as described previously (Huang *et al*., 2014; Huang *et al*., 2021; Shih *et al*., 2020a; Yen *et al*., 2023). The procedures for each assay are outlined briefly below.

#### Open-field test

Mice were individually placed into the center of a transparent plastic box (40 × 40 × 32.5 cm) and allowed to freely explore the box for 10 min. Mouse behaviors in the box were recorded from above and analyzed using the Smart Video Tracking System (Panlab, Barcelona, Spain). Total traveling distance (representing horizontal locomotion activity), number of rearing events (representing exploratory and vertical locomotion activity), number of facial (F) and body (B) grooming events, and the ratio of time spent in the corners to the total time spent in the corners plus central area (representing the degree of anxiety) were measured and analyzed.

#### Elevated plus maze

The apparatus raised to a height of 45.5 cm above the laboratory floor consisted of two open arms (30 cm × 5 cm), and two closed arms (30 cm × 5 cm) enclosed by 14 cm-high walls extending from a central platform (5 cm × 5 cm). An individual mouse was placed on the central area facing the open arm and allowed to freely explore the entire maze for 10 min. Movement was recorded from above and analyzed using the Smart Video Tracking System (Panlab, Barelona, Spain). The percentage of time spent in the open arms, closed arms, and central area were determined. A higher percentage in the closed arm indicates a higher level of anxiety.

#### Three-chamber (3-C) test

A transparent plastic box (17.5 × 41.5 × 22 cm) was divided into three equal compartments by two sliding doors. An identical cylindrical wire cage (16 cm in height, 9.5 cm in diameter) was placed individually in each of the two side chambers. Right before conducting the test, the test mice were placed into a new nesting cage. The entire test consisted of three 10-min sessions, i.e., habituation, sociability, and novelty preference tests, with 2-min intervals between each session. In the first habituation session, both cylindrical wire cages were empty. In the second sociability test, a plastic toy of a cartoon character of similar size to test mice was used as an inanimate object (Ob), which was randomly placed in the wire cage of either the left or right chamber. An unfamiliar mouse of the same sex and of similar size and age (S1) was placed in the other wire cage in the opposite chamber. In the last social novelty session, the inanimate object was replaced by another unfamiliar mouse (S2). The movement and sniffing behaviors of mice were recorded by videotaping from above. Compartments and wire cages were cleaned with 75% EtOH and water after each task. During intervals, the test mice were placed back in the nesting cage. Sniffing toward the cylindrical wire cages was quantified manually without knowing the genotype and treatment of the mice. The total sniffing time was used to represent social interaction. The values of (T_S1_-T_Ob_) and (T_S2_-T_S1_) were determined as preference indices of sociability and social novelty, respectively.

#### Reciprocal social interaction test

The lid of the home cage of test mice was removed to habituate mice for at least 5 min. WT unfamiliar mice of the same sex and similar age and body weight were introduced into the home cage of the test mice. Interactions were video-recorded from above for 5 min. Time spent in social interactions, including sniffing, following and allogrooming, was recorded and analyzed blind to the genotype.

### Quantification and statistical analysis

All images were analyzed using ImageJ. Mouse behavior and neuronal morphology experiments were randomly conducted blindly to avoid personal bias during quantifications. Statistical analysis and graphical outputs were performed using PRISM 10.2.3 (GraphPad software), and the graphics are presented as mean ± SEM and individual data points. All data have been tested for normality (D’Agostino and Shapiro–Wilk tests), unless the sample sizes were unsuitable. To compare two independent groups of data, a two-tailed unpaired *t*-test was used for normally distributed data or a two-tailed Mann–Whitney test was applied for non-normally distributed data. To compare two variants within one dataset, two-way ANOVA with Bonferroni or Tukey multiple comparison test was used. For all comparisons, *P* < 0.05 was considered significant.

## Results

### Body growth of male mice is more sensitive to zinc deprivation than that of female mice

To further evaluate the sensitivity of male and female *Cttnbp2^+/M120I^* mice to dietary zinc, we here subjected male and female *Cttnbp2^+/M120I^* mice and their wild-type littermates to zinc deprivation. We conducted two zinc deprivation protocols, whereby the mice either started on a 150- or 30-ppm zinc diet from the embryonic stage and then both groups shifted to a 0.85-ppm zinc diet (represented as 0-ppm zinc) at postnatal day 28 (P28) until the end of the experiment, i.e., P42, designated as “150-ppm deprivation (150 –> 0)” or “30-ppm deprivation (30 –> 0)”, respectively. The 30-ppm zinc diet represents a “normal” zinc diet, which is sufficient for maintaining the health of wild-type mice for their complete life cycle. Although 150 ppm is much higher than 30 ppm, it is still a reasonable concentration for mice and does not cause side effects (Fourie *et al*., 2018). For each protocol, we compared both male and female mutant mice with their WT littermates of the same sex (**Figure 1A, 1D, 1G, 1J**).

**Figure 1.**
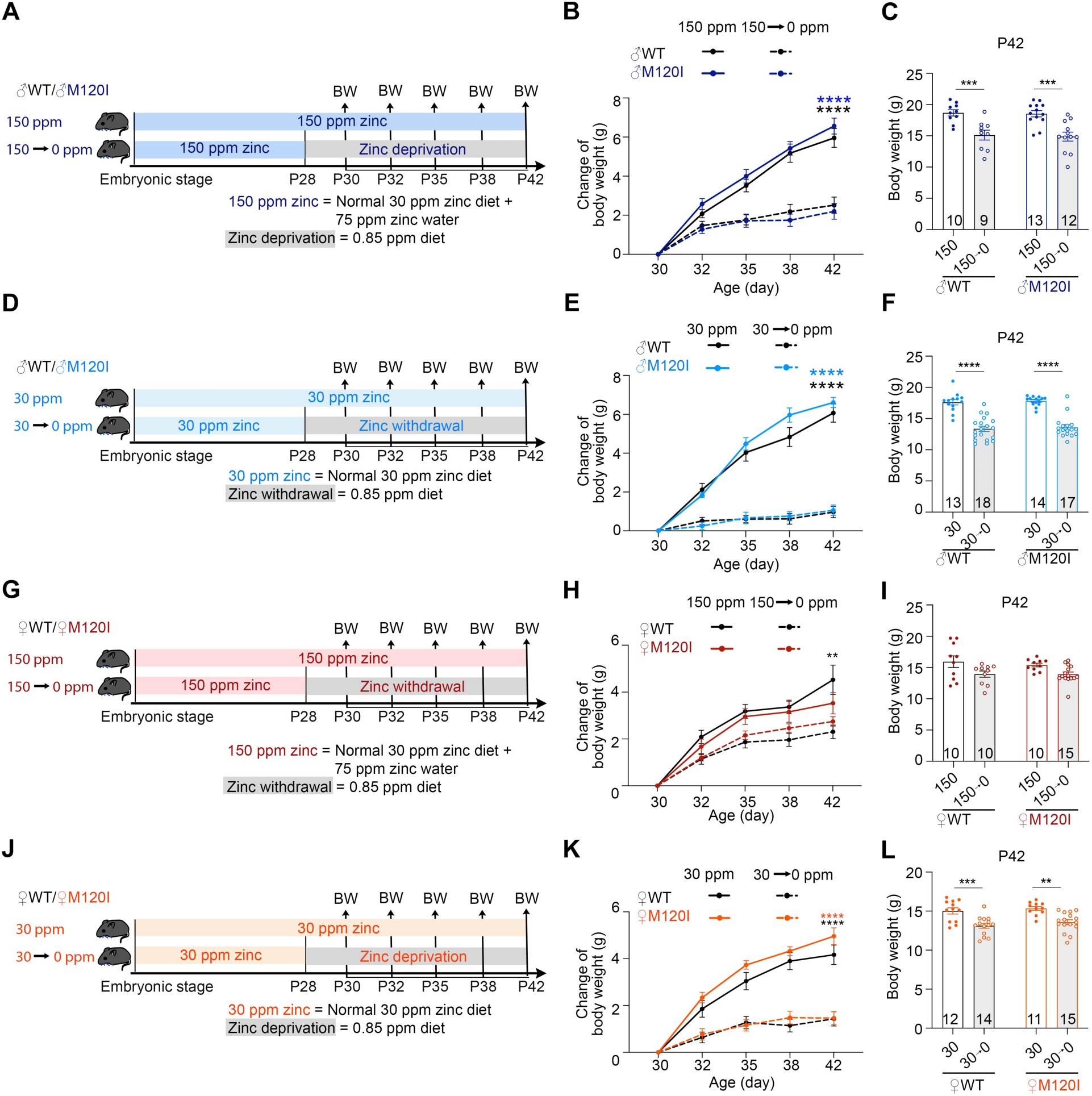
Influence of zinc deprivation on the body weight of both male and female *Cttnbp2^+/M120I^* mice. (**A**) Flow chart of the experimental design to evaluate the effect of 150-ppm deprivation on body weight. Male *Cttnbp2^+/M120I^* mice and WT littermates were fed on 150 ppm zinc diets starting from the embryonic stage. Approximately half of the mice were changed to the 0.85 ppm zinc diet (indicated as 0 ppm in the panel) starting on postnatal day (P) 28, i.e., after weaning. The body weight of all mice was then measured on P30, P32, P35, P38 and P42. (**B**) The results of body weight changes from the experiment in (**A**). (**C**) Average body weight at P42 from (**A**). (**D**) Flow chart of the experimental design of 30-ppm deprivation using male mice. (**E**) The body weight changes from the experiment in (**D**). (**F**) Average body weight at P42 from (**D**). (**G**) Flow chart of the experimental design of 150-ppm deprivation using female mice. (**H**) The body weight changes from the experiment in (**G**). (**I**) Average body weight at P42 from (**G**). (**J**) Flow chart of the experimental design of 30-ppm deprivation using female mice. (**K**) The body weight changes from the experiment in (**J**). (**L**) Average body weight at P42 from (**J**). Sample sizes (N, i.e., mouse numbers) are indicated in (**C**), (**F**), (**I**) and **(L**). Data represent mean values ± SEM. Individual data points are also shown. ** *P* < 0.01; **** *P* < 0.0001; two-tailed unpaired *t* test.

Since zinc is required for cell growth (Brown *et al*., 2002), first we monitored the changes in body weight from P30 to P42 during zinc deprivation (**Figure 1B, 1E, 1H, 1K**), as well as the body weight at the endpoint (P42) of the experiment (**Figure 1C, 1F, 1I, 1L**), to confirm the effect of zinc deprivation. Regardless of genotype, we observed that the body weight of male mice was reduced in both the 30- and 150-ppm deprivation groups compared to the control groups without zinc deprivation (**Figure 1B, 1C, 1E, 1F**). The effects of zinc deprivation were milder for female mice. A clear reduction in female mouse body weight was found only for the 30-ppm deprivation group but not the 150-ppm deprivation group (**Figure 1H-1I, 1K-1L**). Thus, these results indicate that male mice are sensitive to zinc deprivation in terms of maintaining their body growth, whereas female mice appear to be relatively resilient to zinc deprivation, especially when they originally had a higher zinc intake. This observation is similar to humans, with men requiring more zinc to maintain health.

### Only *Cttnbp2^+/M120I^* male mice display impaired social behaviors upon zinc deprivation

Next, we assessed the impact of zinc deprivation on mouse social behaviors. Building on our previous study, we conducted a series of behavioral tests, including open field (OF), elevated plus maze (EPM), three-chamber (3C, which measures sociability and social novelty), and reciprocal social interaction (RSI) test were conducted at postnatal days P30, P32, P35 and P42, respectively, for both male and female mice subjected to 30- and 150-ppm deprivation (**Figure 2A, 2D, 3A, 3D; Supplementary Figure S1A, S1D, S2A, S2D**). In prior studies, we showed that *Cttnbp2* M120I mutation does not affect mouse behaviors in the open field and elevated plus maze test (Shih *et al*., 2020a; Yen *et al*., 2023). However, we included these two tests in our current experiment to maintain consistency with our previous workflow. Similarly, we found no significant differences between the behavior of male and female *Cttnbp2^+/M120I^* mice and their WT littermates in either the open field or elevated plus maze under zinc deprivation (**Supplementary Figures S1-S2**).

**Figure 2.**
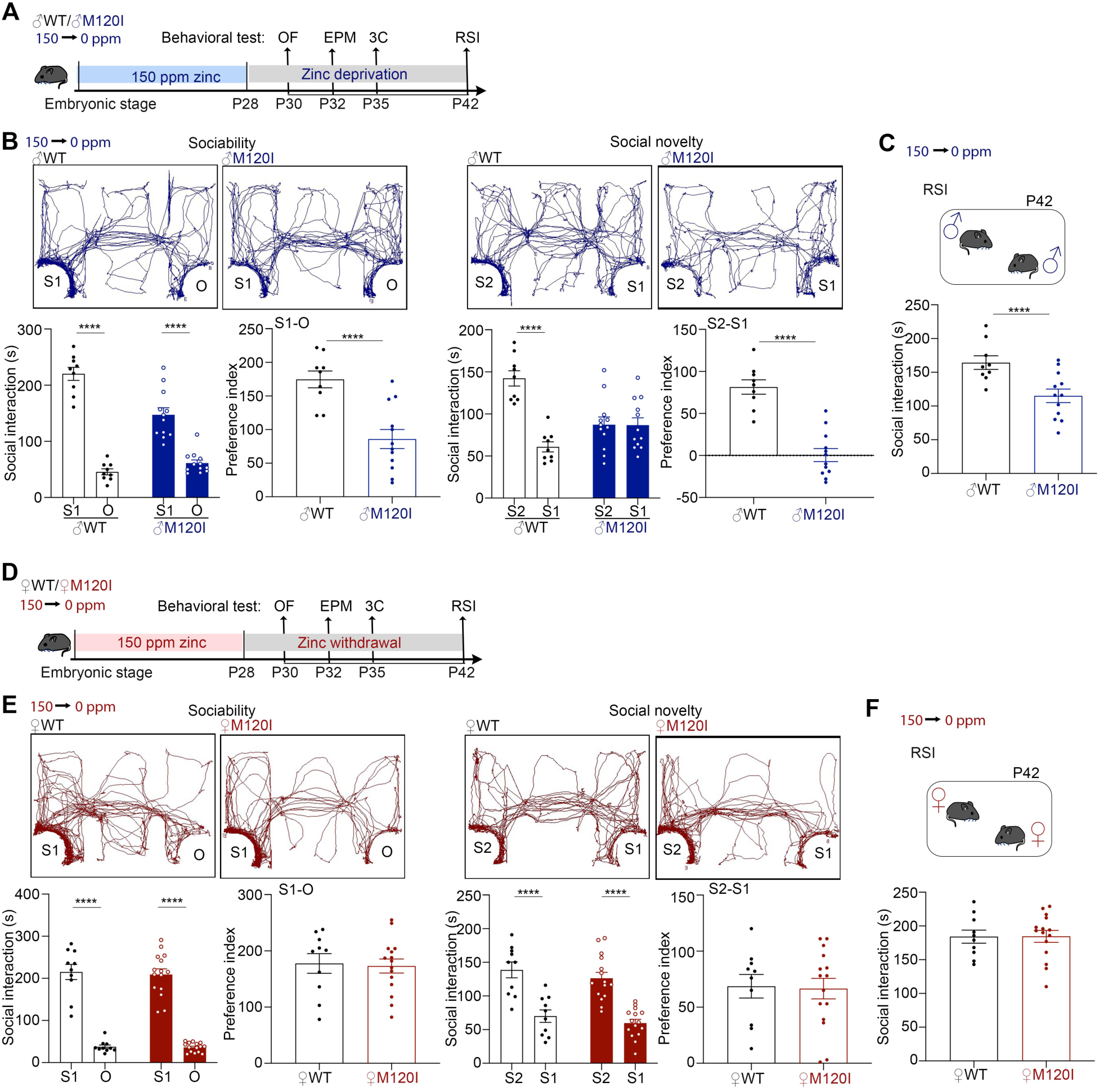
A 150-ppm zinc deprivation diet influences the social behaviors of male *Cttnbp2^+/M120I^* mice, but not those of females. (**A**) Schematic of the experimental design to investigate the effect of 150-ppm zinc deprivation on male mouse behaviors. OF, open field; EPM, elevated plus maze; 3C, three-chamber test; RSI, reciprocal social interaction. (**B**) Results of the three-chamber test from the experiment in (**A**). (**C**) Results of reciprocal social interaction from (**A**). (**D**) Schematic of the experimental design to investigate the effect of 150-ppm zinc deprivation on female mouse behaviors. (**E**) Results of the three-chamber test from the experiment in (**D**). (**F**) Results of reciprocal social interaction from (**D**). Data represent mean values ± SEM. Individual data points are also shown. **** *P* < 0.0001; two-tailed unpaired *t* test, except for the three-chamber test for which interaction times were analyzed by two-way ANOVA with Tukey’s multiple comparisons post-hoc test.

**Figure 3.**
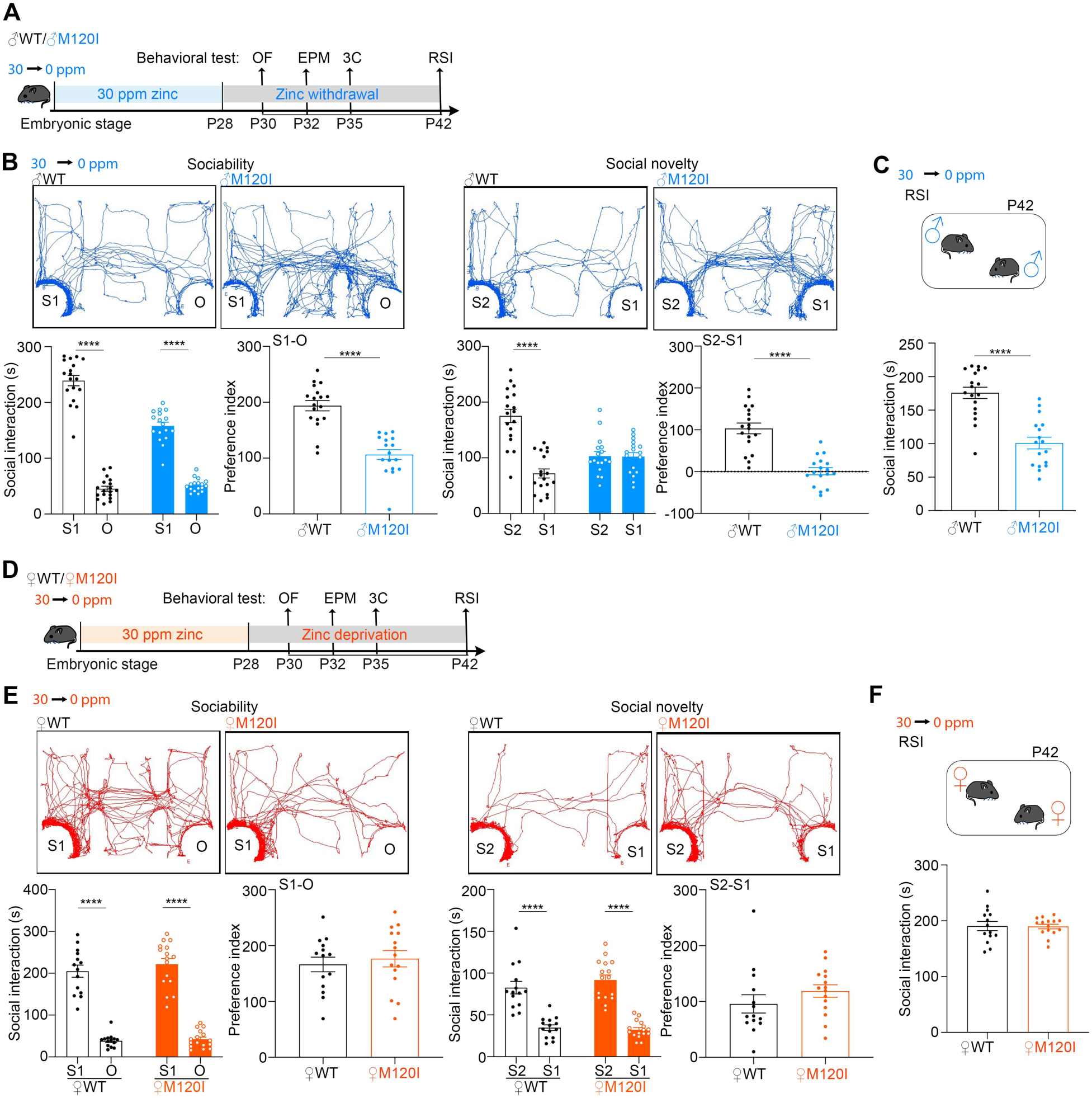
Female *Cttnbp2^+/M120I^*mice are resilient to 30-ppm zinc deprivation in terms of social behaviors. (**A**) Schematic of the experimental design to investigate the effect of 30-ppm zinc deprivation on male mouse behaviors. OF, open field; EPM, elevated plus maze; 3C, three-chamber test; RSI, reciprocal social interaction. (**B**) Results of the three-chamber test from the experiment in (**A**). (**C**) Results of reciprocal social interaction from (**A**). (**D**) Schematic of the experimental design to investigate the effect of 30-ppm zinc deprivation on female mouse behaviors. (**E**) Results of the three-chamber test from the experiment in (**D**). (**F**) Results of reciprocal social interaction from (**D**). Data represent mean values ± SEM. Individual data points are also shown. **** *P* < 0.0001; two-tailed unpaired *t* test, except for the three-chamber test for which interaction times were analyzed by two-way ANOVA with Tukey’s multiple comparisons post-hoc test.

Our previous studies have shown that autism-linked behaviors associated with *Cttnbp2* deficiency are present only in male mice, but not in female mice, when fed a 30 or 84 ppm zinc diet, compared to their wild-type (WT) littermates of the same sex (Shih *et al*., 2022; Shih *et al*., 2020a; Shih *et al*., 2020b; Yen *et al*., 2023). Increasing the dietary zinc level to 150 ppm improved the social behaviors of male Cttnbp2-deficient mice (Shih *et al*., 2022; Shih *et al*., 2020a; Yen *et al*., 2023). However, when subjected to 150-ppm zinc deprivation, male *Cttnbp2^+/M120I^*mice exhibited social deficits (**Figure 2A-2C**). For male WT mice, no matter whether constantly feeding 150 ppm zinc diet (Yen *et al*., 2023) or undergoing 150-ppm zinc deprivation (**Figure 2C**), male WT mice spent similar amounts of time (∼160 sec) interacting with stranger mice in RSI. It suggests that male WT mice are resilient to two weeks of zinc deprivation. However, 150-ppm deprivation reduced the interaction of male *Cttnbp2^+/M120I^* mice with stranger S1 in the sociability test and lowered the preference for the S2 stranger in the social novelty test of the three-chamber test (**Figure 2B**). In the RSI test, 150-ppm deprivation also resulted in a shorter interaction time between male *Cttnbp2^+/M120I^* mice and unfamiliar mice compared to male WT mice undergoing the same treatment (**Figure 2C**).

In contrast, female *Cttnbp2* mutant mice behaved comparably to their WT littermates when they were subjected to 150-ppm deprivation. In both the three-chamber and reciprocal social interaction tests, the performances of female *Cttnbp2^+/M120I^* mice were equivalent to those of their WT littermates (**Figure 2E-2F**). Thus, 150-ppm zinc deprivation does not appear to have a notable impact on the behavioral repertoire of female *Cttnbp2* mutant mice relative to WT counterparts.

In terms of the 30-ppm deprivation group, an experimental condition in which the mice were raised with a much-reduced zinc intake before being subjected to further deprivation compared to the 150-ppm deprivation group, as expected the male *Cttnbp2^+/M120I^* mice exhibited social deficits (**Figure 3A-3C**). Importantly, female *Cttnbp2^+/M120I^* mice of the 30-ppm deprivation group still appeared to behave normally, with their performance in the three chamber and receiprocal social interaction tests being comparable to their WT littermates (**Figure 3D-3F**). In addition, the performances of female mice upon 150- or 30-ppm deprivation were found previously to be comparable to those fed continuously on a 150- or 30-ppm zinc diet (Yen *et al*., 2023), further supporting the insensitivity of female mice to zinc deprivation. These outcomes indicate that even though body growth is restricted by 30-ppm zinc deprivation, the social behaviors of female *Cttnbp2^+/M120I^*mice remain unaffected.

Taken together, our behavioral assays reveal that male *Cttnbp2^+/M120I^* mice, but not female mice, are sensitive to zinc deprivation. Thus, ASD-linked *Cttnbp2* mutation and male sex render mice susceptible to zinc deficiency.

### Zinc is highly relevant to ASD-linked genes

To further explore the role of zinc in the sex bias of ASD, we first analyzed all 1191 ASD-linked genes collected by the Simons Foundation Autism Research Initiative (SFARI, https://www.sfari.org/resource/sfari-gene/, August 19, 2024 release) and identified 220 zinc-related ASD genes, accounting for more than 18% of SFARI genes (**Figure 4A, 4B, Supplementary Table S1**). STRING network analysis revealed that 199 of the zinc-related SFARI proteins bind metal ions, 156 are zinc finger proteins (accounting for 13% of SFARI genes), and 73 are DNA-binding (**Figure 4A**). Given that 3% of all human genes encode zinc finger proteins (Klug, 2010), it is clear that zinc finger proteins are enriched among SFARI genes (up to 13%). Regarding biological processes, most of the zinc finger proteins are involved in transcription (107 genes) and metabolic processes (136 genes), with some others related to neuronal differentiation (10 genes) and neurogenesis (29 genes) (**Figure 4B**).

**Figure 4.**
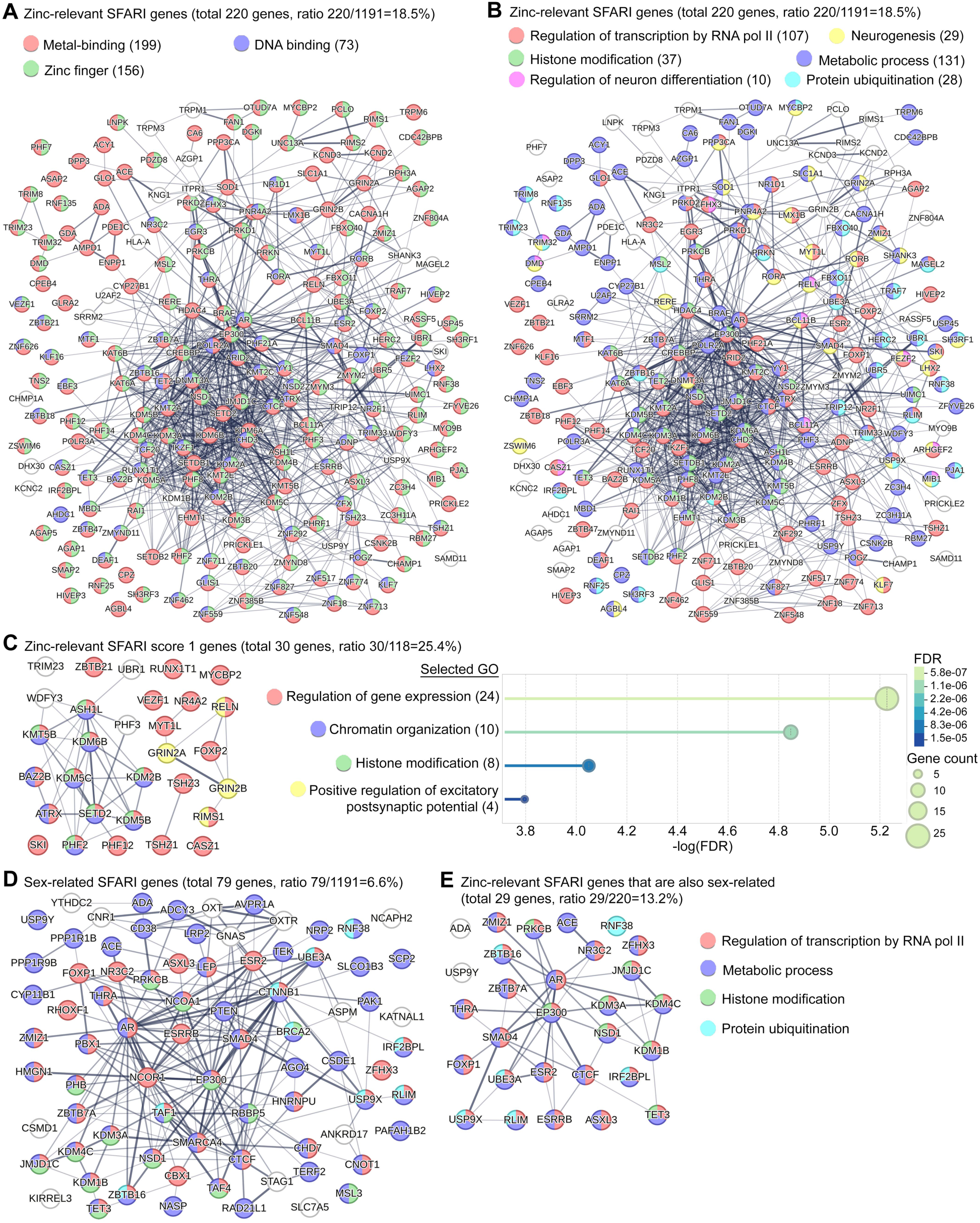
Protein networks of zinc- and sex-related SFARI genes analyzed using STRING. (**A**) Molecular functions of 220 zinc-related SFARI genes. (**B**) Biological processes of 220 zinc-related SFARI genes. (**C**) Biological processes of 30 zinc-related SFARI Score 1 genes. Some significant Go are summarized in the right panel. The false discovery rate and gene counts of the GO are also indicated. (**D**) Biological processes of 79 SFARI genes related to at least one keyword from among estrogen, androgen, sex, male, female, and thyroxine. (**E**) Biological processes of 29 zinc-related SFARI genes that are also related to at least one keyword from among estrogen, androgen, sex, male, female, and thyroxine. Different colored balls represent the different GO terms, as indicated. The numbers in the brackets are the total numbers of genes belonging to those GO terms. In (**D**) and (**E**), the same colors are used to represent the indicated biological processes. The complete gene lists are presented in **Supplementary Tables S1-S3**.

When analyzing 118 SFARI Score 1 genes, excluding syndromic genes, we identified 30 zinc-related genes (**Figure 4C**). These account for over 25% of Score 1 genes, surpassing the ∼18% seen in all SFARI genes. Since Score 1 genes meet the strictest criteria for ASD causative genes, this enrichment further supports the link between zinc and ASD. Like other zinc-related SFARI genes, zinc finger proteins dominate, with 23 genes in this group. Key GO terms include gene expression regulation, chromatin organization, and histone modification. Additionally, four genes, including *GRIN2A* and *GRIN2B*, contribute to excitatory postsynaptic potential regulation (**Figure 4C**). These network analyses strengthen the association between zinc and ASD, suggesting that zinc influences neuronal processes, particularly through transcriptional and metabolic regulation.

Next, we searched for potential sex-relevant genes from among the zinc-related SFARI genes and total SFARI genes using the keywords “estrogen”, “androgen”, “sex”, “male”, “female” and “thyroxine” in UniProt (https://www.uniprot.org). From among the 1191 SFARI genes, we identified 79 as potentially being sex-relevant, accounting for 6.6% of all SFARI genes (**Figure 4D, Supplementary Table S2**). In terms of zinc-related SFARI genes, 29 were potentially linked to a sex bias response, accounting for 13% of the zinc-related SFARI genes (**Figure 4E, Supplementary Table S3**). Moreover, 25 of the 29 genes were associated with transcriptional regulation (**Figure 4E**). These analyses support a role for transcription in the zinc-related sex bias of ASD.

### Long-term dietary zinc supplementation promotes ribosome biogenesis and the metabolic processes of male *Cttnbp2^+/M120I^* mice

In addition to the effects of zinc on transcription, as described above, we further investigated how dietary zinc supplementation differentially influences the synapses of male and female *Cttnbp2^+/M120I^* mice. First, we examined the beneficial effects of a 150-ppm zinc diet on male mice. Synaptosomal fractions prepared from male mice fed on 150 ppm or 30 ppm zinc diets starting from the embryonic stage were subjected to liquid chromatography with tandem mass spectrometry (LC-MS-MS). We cross-compared the four groups of samples, i.e., male WT and *Cttnbp2^+/M120I^* mice with a 30 or 150 ppm zinc diet (designated as 30WT, 150WT, 30MI and 150MI), to evaluate the genetic and dietary effects on male brains (**Figure 5, Supplementary Figure S3, S4 and Tables S4-S5**). The differentially expressed proteins (DEPs) were selected based on an adjusted *p*-value of < 0.05 and reliable peptide signals.

**Figure 5.**
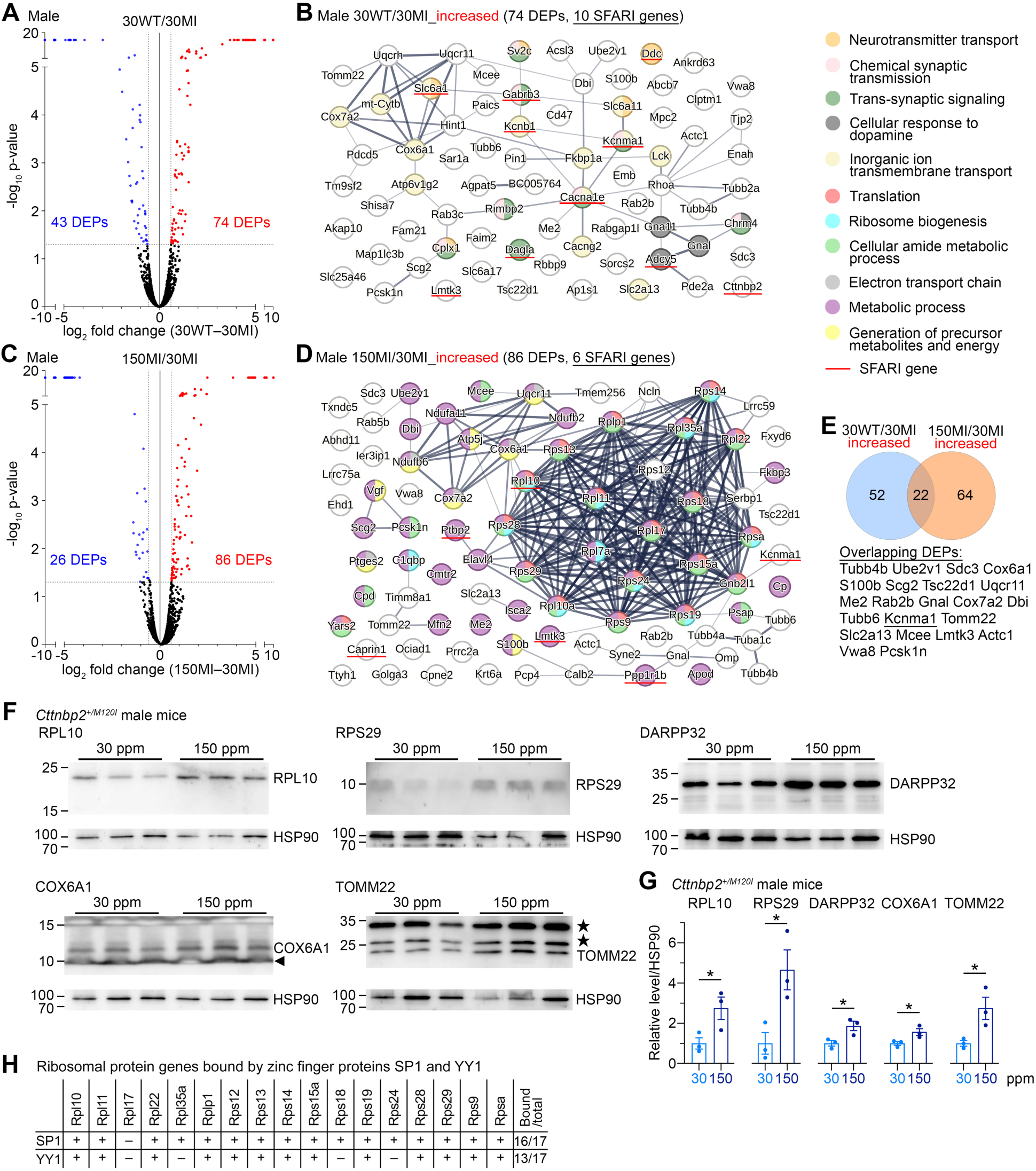
Long-term zinc supplementation alters the synaptic proteomes of male *Cttnbp2^+/M120I^* mice. Male *Cttnbp2^+/M120I^* mice and wild-type littermates fed on either a 30 ppm or 150 ppm zinc diet starting from the embryonic stage were subjected to synaptic proteome analysis at P42. Comparisons of WT mice fed the 30 ppm zinc diet (30WT) vs. *Cttnbp2^+/M120I^* mice fed on the 30 ppm zinc diet (30MI), as well as the *Cttnbp2^+/M120I^* mice fed the 150 ppm zinc diet (150MI) vs. those on the 30 ppm zinc diet (30MI) were performed. (**A**) Volcano plot of the 30WT vs. 30MI comparison. Numbers of differentially expressed proteins (DEPs) displaying increased or decreased expression are indicated. (**B** STRING network of increased DEPs from the 30WT vs. 30MI comparison. (**C**) Volcano plot of the 150MI vs. 30MI comparison. Numbers of DEPs displaying increased or decreased expression are indicated. (**D**) STRING network of increased DEPs from the 150MI vs. 30MI comparison. (**E**) Increased DEPs shared between the 30WT/30MI and 150MI/30MI comparative groups. (**F**) Immunoblots of RPL10, RPS29, DARPP32, COX6A1 and TOMM22, validating the proteomics results. (**G**) Quantification of (**F**). HSP90 was used as a loading control. The sample size (N = 3) indicates the number of mice subjected to analysis. Data represent mean values ± SEM. Individual data points are also shown. *, *P* < 0.05; two-tailed unpaired *t* test. (**H**) Summary of SP1 and YY1 binding to ribosomal protein genes. The complete lists of DEPs are available in **Supplementary Tables S4 and S5**. The GO terms and SynGO analysis of increased DEPs of the 30WT vs. 30MI and 150MI vs. 30MI comparisons are shown in **Supplementary Figure S3**. The 150WT vs. 150MI and 150WT vs. 30WT comparisons are summarized in **Supplementary Figure S4**. The full-size uncropped blots of (**F**) are shown in **Supplementary Figure S5**.

The 30WT and 30MI groups were first compared to reveal the effect of *Cttnbp2* mutation under the condition of the 30-ppm zinc diet, which uncovered 117 DEPs (**Figure 5A**, 74 and 43 DEPs displaying increased or decreased expression levels, respectively). The major gene ontology (GO) terms of all those DEPs are highly relevant to synapses (**Supplementary Figure S3A**), consistent with our previous results that Cttnbp2 deficiency impairs synaptic organization (Shih *et al*., 2020a; Yen *et al*., 2023). Among the 74 increased DEPs, 24 are SynGO proteins (Synaptic Gene Ontologies, https://www.syngoportal.org), which were enriched in post- and pre-synaptic processes, especially in synaptic vesicle exocytosis (**Supplementary Figure S3B**). STRING analysis (https://string-db.org) also indicated neurotransmitter transport, chemical synaptic transmission, trans-synaptic signaling, cellular response to dopamine, and inorganic ion transmembrane transport as being significant GO terms (**Figure 5B**). Among these increased DEPs, 10 are also SFARI genes (https://gene.sfari.org/database/human-gene/) (**Figure 5B**, underlined). The altered expression of these ASD- and synapse-related proteins may contribute to the synaptic phenotypes and ASD-linked behaviors of male *Cttnbp2^+/M120I^*mice having a 30 ppm zinc diet.

The 150MI/30MI comparison that investigated the beneficial effect of 150 ppm zinc supplementation identified 112 DEPs (**Figure 5C**, 86 increased and 26 decreased DEPs). In contrast to the results of the 30WT/30MI comparison, we noticed that translation, ribosome biogenesis, metabolic process, generation of precursor metabolites, and energy and electron transport chain were the major GO terms (**Supplementary Figure S3C**). Thirty-two of the DEPs exhibiting increased expression levels were identified as SynGO proteins. Metabolism-associated GO terms, especially involving synaptic ribosomes, were highly enriched among the 86 increased DEPs of the 150MI/30MI comparison (**Supplementary Figure S3D**). STRING analysis further highlighted that the protein expression levels of 19 ribosomal proteins, 7 mitochondrial proteins, and many others involved in metabolic processes, were increased in the brains of the 150MI group (**Figure 5D**). These results indicate the significance of ribosome biogenesis and metabolic processes in the response to long-term zinc supplementation in the male mouse brain.

Only 22 DEPs (< 1/3 of DEPs) were shared between the 30WT/30MI increased and 150MI/30MI increased comparisons (**Figure 5E**). Among these shared DEPs, only KCNMA1 is relevant to synaptic function (**Figure 5E**). Thus, long-term zinc supplementation may not directly increase levels of proteins involved in neurotransmission or synaptic function, but may increase protein synthesis and metabolic processes to indirectly ameliorate the deficits of synapse formation and function caused by *Cttnbp2* deficiency.

For the comparisons of the male 150WT/150MI and 150WT/30WT groups, we also identified several synaptic and ribosomal proteins, respectively, supporting the influence of *Cttnbp2* deficiency and dietary zinc on synaptic proteomes (**Supplementary Figure S4**).

To validate the results of our LC-MS-MS and bioinformatic analyses, we picked several of the increased DEPs from the 150MI/30MI comparison for immunoblotting, including the ribosomal proteins RPL10 and RPS29, two mitochondrial proteins TOMM22 and COX6A1, and the signaling molecule DARPP32. The immunoblotting results confirmed that expression levels of all five of these proteins were indeed increased in male *Cttnbp2^+/M120I^*mice by long-term 150 ppm zinc supplementation (**Figure 5F-5G, Supplementary Figure S5**), supporting the reliability of our proteomic analysis.

### Potential for transcriptional control to be involved in zinc-regulated ribosomal biogenesis

We detected considerable upregulation of many ribosomal proteins in response to zinc supplementation in male *Cttnbp2^+/M120I^*mice (**Figure 5D**). Previous studies have indicated that mammalian ribosomal proteins are controlled by common transcriptional factors, including SP1, YY1 and GABP (Perry, 2005; Petibon *et al*., 2021), to ensure relatively equimolar expression of ribosomal proteins. Among these three transcriptional factors, SP1 and YY1 are both zinc finger proteins and their activities are modulated by zinc (Figiel *et al*., 2023a; Figiel *et al*., 2023b; Torigoe *et al*., 2003). Thus, dietary zinc supplementation may regulate ribosome biogenesis via transcriptional regulation of zinc finger proteins. To evaluate that possibility, we searched an existing chromatin-immunoprecipitation database (Vorontsov *et al*., 2018) to confirm if SP1 and YY1 bind to the 19 ribosomal protein genes shown in **Figure 5D**. Since reliable data were not available for Rpl10a and Rpl17a (Vorontsov *et al*., 2018), we analyzed the remaining 17 (**Figure 5H**). We found that SP1 and YY1 bound 16 and 13 of those 17 ribosomal protein genes, respectively (**Figure 5H**). The only ribosomal protein gene that did not host either an SP1 or YY1 site was *Rpl17* (**Figure 5H**). However, the *Rpl17* gene contains several binding sites for other zinc finger transcription factors, including members of the ZNF and KLF families (https://genome.ucsc.edu/cgi-bin/hgSearch?search=Rpl17&db=mm9). Therefore, expression of the *Rpl17* gene is likely controlled by other zinc finger proteins. These bioinformatic analyses support that the enhancement of ribosome biogenesis by zinc in male *Cttnbp2^+/M120I^* mutant brains likely acts through zinc finger transcription factors.

### Translation-related regulation in the female brain is affected by *Cttnbp2* mutation

To investigate how female *Cttnbp2^+/M120I^*mice are resilient to zinc deprivation, we analyzed the DEPs in the synaptosomal proteomes of female mice first fed on a 30-ppm zinc diet and then subjected to 30-ppm zinc deprivation. We focused on three comparative groupings, i.e., 30WT/30MI, 0WT/0MI and 30MI/0MI (**Figure 6A, Supplementary Table S6-S8**), where the “0” here indicates 30-ppm deprivation. The GO of these DEPs proved less informative than for our male data (**Figure 6B-6E**). However, we noticed that several proteins involved in translation-related regulation were present in the decreased DEPs of the 30WT/30MI and/or 0WT/0MI comparisons. Notably, the translation initiation factors EIF1 and EIF1A were identified among the decreased DEPs of both comparative groups (**Figure 6A-6C**). The results of immunoblotting confirmed higher protein levels of EIF1 and EIF1A in female *Cttnbp2^+/M120I^*mice compared to their WT littermates (**Figure 6F**). The higher expression levels of EIF1 and EIF1A may promote translation in female mice. This observation is consistent with findings from our previous study showing that female *Cttnbp2^+/M120I^* mice fed on an 84-ppm zinc diet exhibit greater mTOR activity (Yen *et al*., 2023), a critical kinase responsible for controlling protein synthesis via multiple pathways. Consequently, we also assessed levels of mTOR phosphorylation in the brain of female WT and *Cttnbp2^+/M120I^*mice treated with or without zinc deprivation. We found that mTOR phosphorylation levels were increased in the female *Cttnbp2^+/M120I^*mouse brains after zinc deprivation, but not so in WT females (**Figure 6G**). Thus, the brains of female *Cttnbp2^+/M120I^* mice may be resistant to zinc deprivation by expressing more translation initiation factors and maintaining higher activation of the mTOR pathway, ultimately compensating for the deficits caused by *Cttnbp2* mutation.

**Figure 6.**
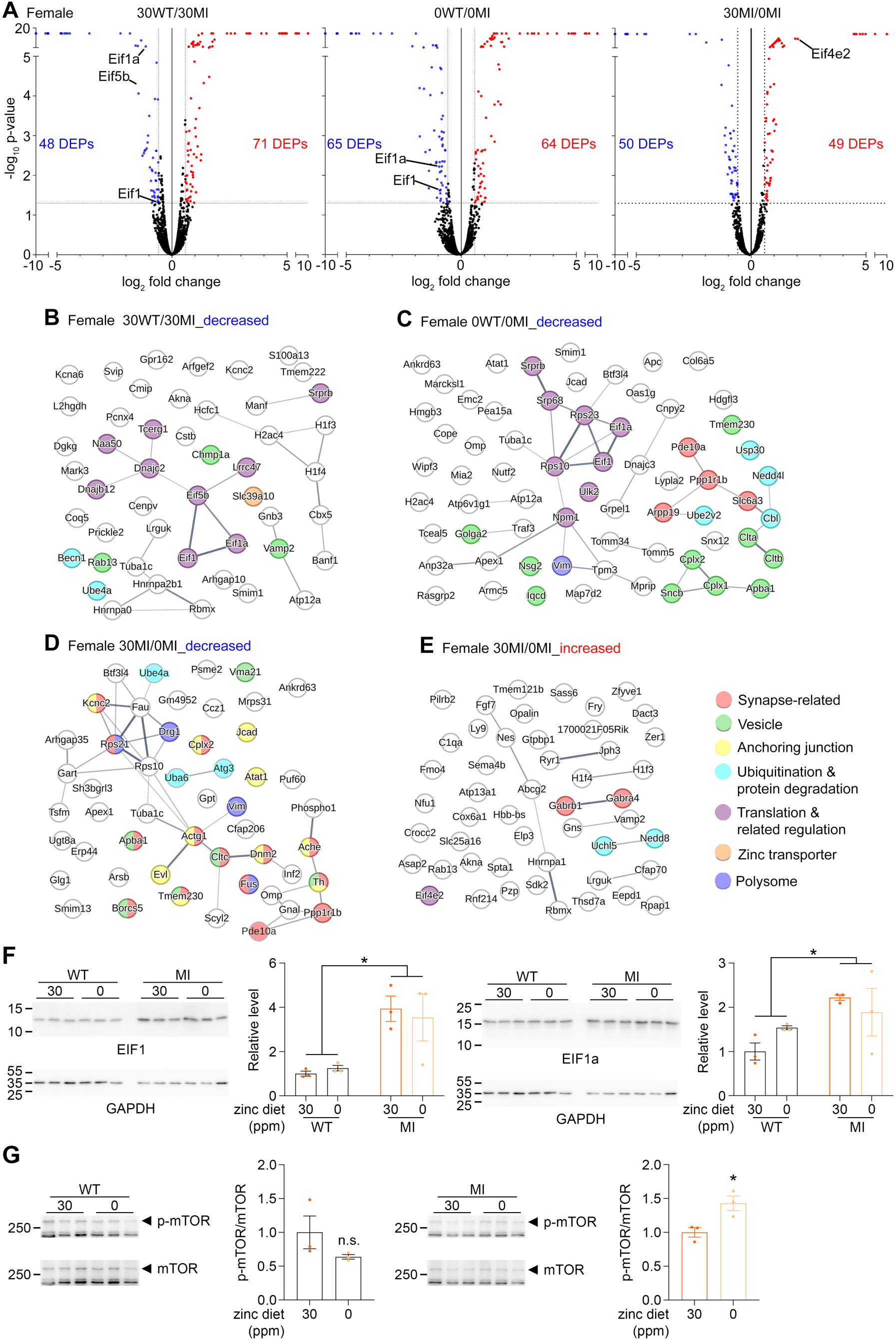
Two-week zinc deprivation alters the synaptic proteomes of female *Cttnbp2^+/M120I^* mice. Female *Cttnbp2^+/M120I^* mice and wild-type littermates were fed on a 30 ppm zinc diet starting from the embryonic stage and then changed to the 0.85 ppm zinc diet at P28, before being subjected to synaptic proteome analysis at P42. Comparisons of WT mice fed on the 30 ppm zinc diet (30WT) vs. *Cttnbp2^+/M120I^* mice fed the 30 ppm zinc diet (30MI), WT mice subjected to zinc deprivation (0WT) vs. *Cttnbp2^+/M120I^* mice with zinc deprivation (0MI), and *Cttnbp2^+/M120I^*mice fed the 30 ppm zinc diet (30MI) vs. those subjected to zinc deprivation (0MI) were performed. (**A**) Volcano plot of the 30WT vs. 30MI, 0WT vs. 0MI, and 30MI vs. 0MI comparison. Numbers of DEPs displaying increased or decreased expression are indicated. Some initiation factors are also shown in the figure. (**B**) STRING network of decreased DEPs for the 30WT vs. 30MI comparison. (**C**) STRING network of decreased DEPs for the 0WT vs. 0MI comparison. (**D**) STING network of decreased DEPs for the 30MI vs. 0MI comparison. (**E**) STRING network of increased DEPs for the 30MI vs. 0MI comparison. (**F**) Immunoblots and quantification results for EIF1 and EIF1A. Three mice for each group were analyzed. Data represent mean values ± SEM. Individual data points are also shown. *, *P* < 0.05; two-way ANOVA. (**G**) Immunoblots and p-mTOR/mTOR ratio quantification. Three mice for each group were analyzed. Data represent mean values ± SEM. Individual data points are also shown. n.s., no significance; *, *P* < 0.05; two-tailed unpaired *t*-test. The complete lists of DEPs are available in **Supplementary Tables S6-8**. The full-size uncropped blots are shown in **Supplementary Figure S6**.

### Long-term zinc supplementation promotes protein synthesis in neuronal cultures

Next, we used cultured neurons to investigate if an increase in zinc concentration indeed promotes their synthesis of proteins (**Figure 4A**). Neurobasal, our culture medium for neuronal cultures, contains only 0.6 μM zinc (https://www.thermofisher.com/tw/zt/home/technical-resources/media-formulation.251.html). An additional 100 μM zinc was included in the culture medium when neurons were plated, with an absence of this extra zinc (labeled as 0 μM) being used as a control. To ensure protein synthesis was not limited by the availability of free amino acids in the culture, amino acid supplements were also added (indicated as extra 1X a.a. in the figures) for both the 0 or 100 μM zinc conditions.

We used Coomassie blue staining and puromycin pulse labeling for 10 min to monitor total protein levels and newly synthesized proteins, respectively (**Figure 7A**). We loaded equal fractions of neuronal lysates collected from individual wells into SDS-PAGE to investigate if zinc treatment for 10 or 18 days increased total protein amounts in the cultures. The results of Coomassie blue staining showed that 100 μM zinc treatment for both 10 and 18 days increased the total protein amounts of neuronal cultures prepared from both WT and *Cttnbp2^+/M120I^*mice (**Figure 7B**, **7C**). Among all comparisons, MI/100 μM vs. MI/0 μM at 18 days in vitro (DIV) displayed the highest increment in total protein (∼1.8-fold) (**Figure 7B, 7C**). Here we used mixed-sex cultures and did not separate male and female neurons. Although the presence of female neurons may dilute the effect of zinc, we still found the differences between 100 vs. 0 μM zinc, supporting the notion that *Cttnbp2^+/M120I^*neurons respond strongly to zinc treatment in terms of protein synthesis.

**Figure 7.**
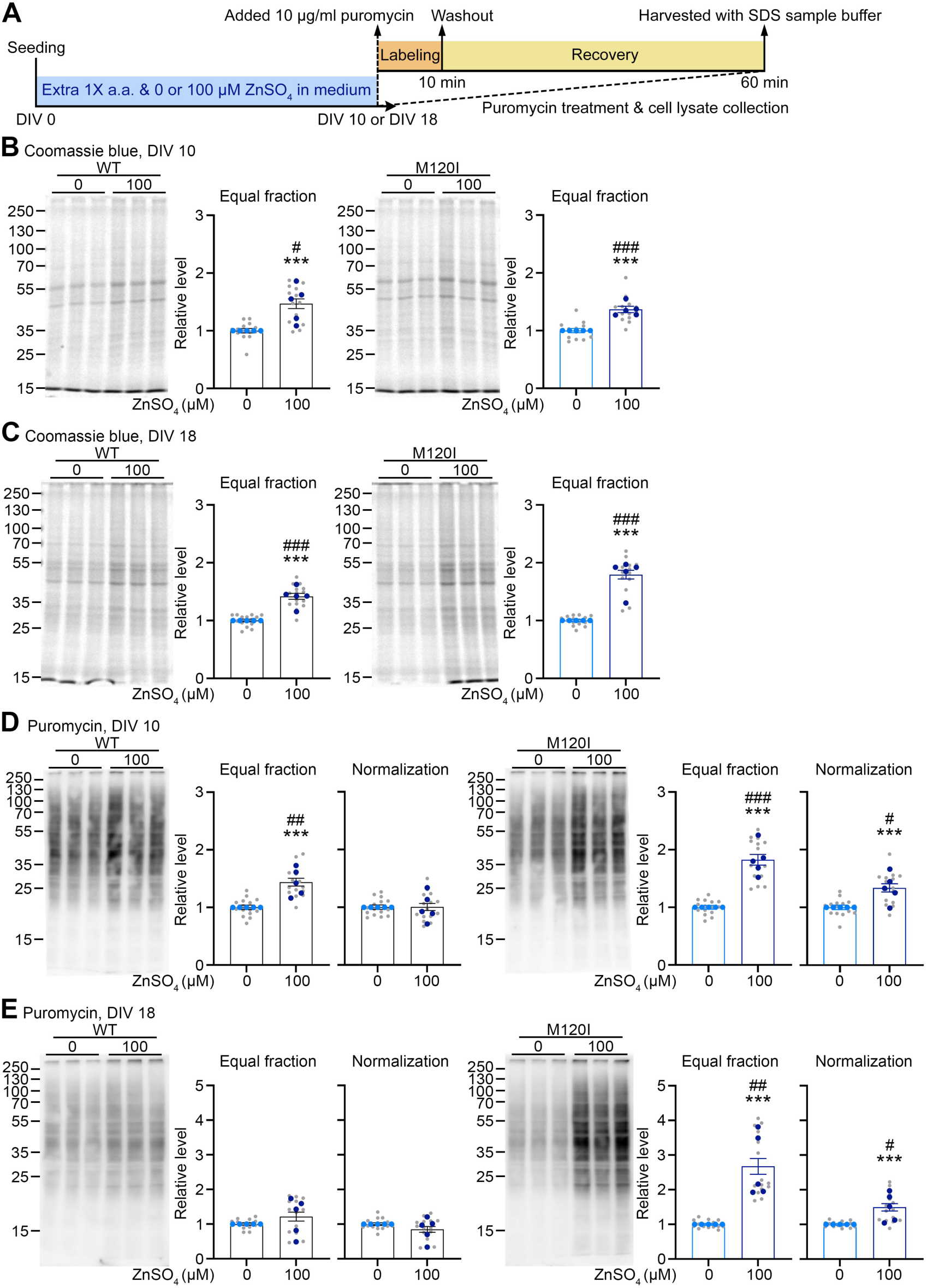
Long-term zinc treatment enhances protein synthesis of neurons cultured *in vitro*. (**A**) Experimental timeline of long-term zinc treatment and puromycin labeling. (**B**) Immunoblots and quantification of Coomassie blue staining at DIV 10. (**C**) Immunoblots and quantification of Coomassie blue staining at DIV 18. (**D**) Immunoblots and quantification of puromycin labeling at DIV 10. (**E**) Immunoblots and quantification of puromycin labeling at DIV 18. (**B-E**) Data represent mean values ± SEM. The gray dots indicate individual samples (N = 15): ***, *P* < 0.001; Two-tailed unpaired *t*-tests were used, except that the result of WT Coomassie blue staining was analyzed using a Mann-Whitney test. The light and dark blue dots indicate the averages of each batch of samples (N = 5); #, *P* < 0.05; ##, *P* < 0.01; ###, *P* < 0.001. In (**D**) and (**F**), normalization indicates that the quantitative results of equal fractions were further normalized against the Coomassie Blue staining signals in (**B**) and (**C**), respectively.

Then, we used two methods to analyze the puromycin pulse labeling results. One was to directly compare the puromycin signals in equal fractions of lysates to reveal the total amounts of proteins synthesized *de novo* in each well. The other was to further normalize the data against the respective Coomassie blue staining signals, thereby reflecting protein synthesis efficiency per unit protein mass (**Figure 7D, 7E**). In our equal fraction analysis, WT neurons displayed stronger puromycin signal under the condition of 100 µM zinc treatment at 10 DIV but not 18 DIV (**Figure 7D, 7E, right**). However, *Cttnbp2^+/M120I^* neurons exhibited stronger puromycin signals at both 10 and 18 DIV in the presence of the extra 100 μM zinc (**Figure 7D, 7E, right**). After normalization against Coomassie blue staining signals, only *Cttnbp2^+/M120I^*neurons displayed higher protein synthesis efficiency per protein mass at 10 and 18 DIV (**Figure 7D, 7E**). Thus, *Cttnbp2^+/M120I^* neurons indeed present a stronger response to zinc treatment in terms of the upregulation of protein synthesis.

Next, we investigated if reducing the timeframe of zinc treatment could have the same effect. To do so, we added 100 μM zinc to neuronal culture at 16 DIV. Two days later, the samples were subjected to Coomassie blue staining and puromycin pulse labeling as described above (**Supplementary Figure S6A**). We found that there were no differences in Coomassie blue staining or puromycin signals between the 0- and 100-μM zinc groups in this set of experiments (**Supplementary Figure 6B, 6C**). Thus, the enhancement of protein synthesis elicited by zinc treatment is a chronic response that requires more than two days of treatment.

We also investigated expression levels of the RPL10 and RPS29 proteins upon zinc treatment of cultured neurons. We observed that, compared to the 0 µM zinc group, the 100 μM zinc treatment increased the protein levels of RPS29 and RPL10 in both WT and *Cttnbp2^+/M120I^* neurons in an equal fraction analysis (**Figure 8A**-**8D**). After normalization against Coomassie blue signals, only RPS29 displayed higher protein synthesis efficiency in *Cttnbp2^+/M120I^* neurons at both 10 and 18 DIV (**Figure 8A, 8B**). These results indicate that although our zinc treatment generally enhanced the protein synthesis efficiency of *Cttnbp2^+/M120I^*neurons, certain proteins, such as RPS29, responded more strongly than others. Nevertheless, these results are consistent with the notion that zinc treatment for 10 or 18 days promotes ribosome biogenesis and protein synthesis of *Cttnbp2^+/M120I^* cultures.

**Figure 8.**
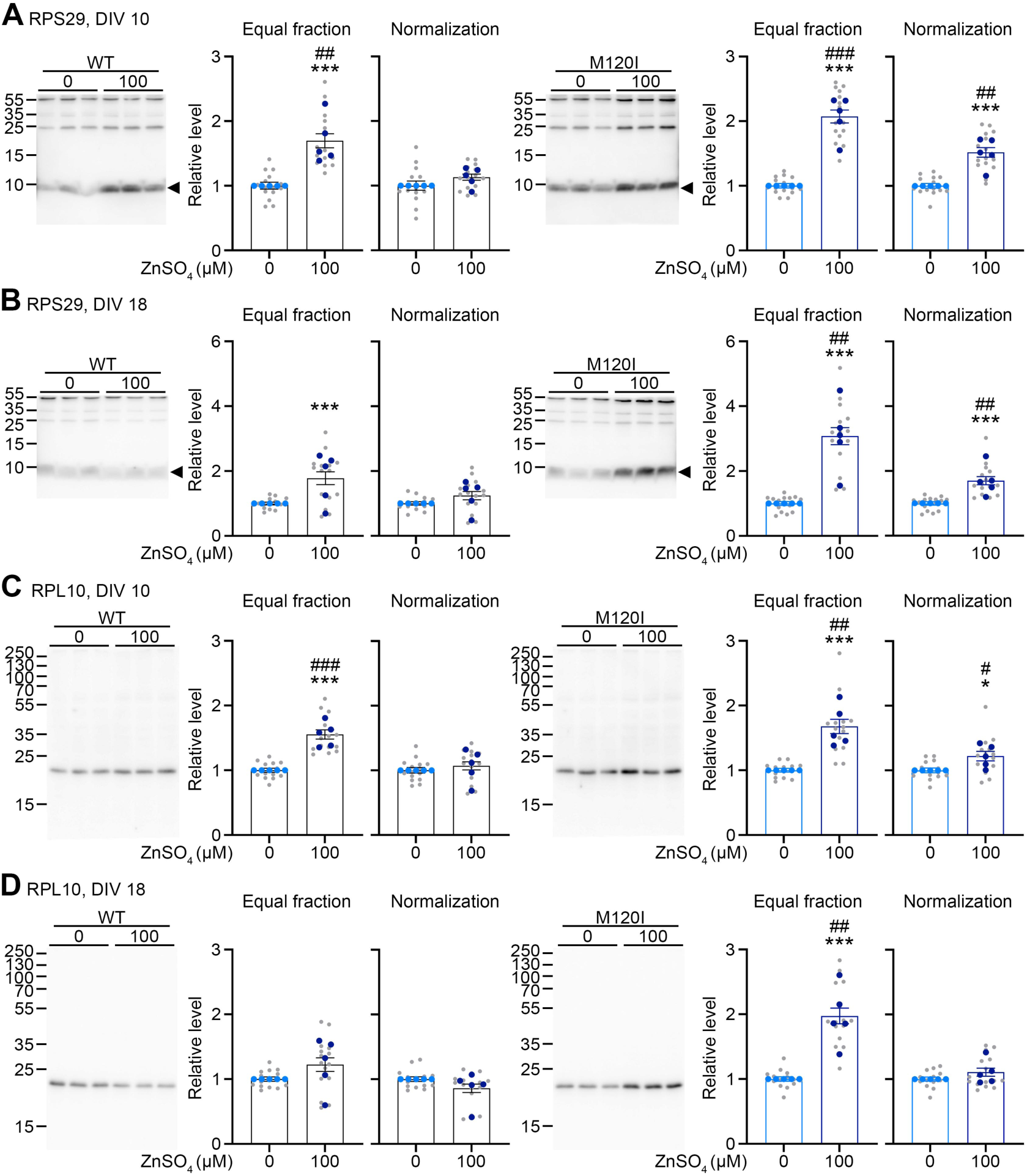
Long-term zinc treatment enhances ribosomal protein expression in culture. (**A**) Immunoblots and quantification of RPS29 at DIV 10. The arrow points to the position of RPS29. (**B**) Immunoblots and quantification of RPS29 at DIV 18. The arrow points to the position of RPS29. (**C**) Immunoblots and quantification of PRL10 at DIV 10. (**D**) Immunoblots and quantification of RPL10 at DIV 18. (**A-D**) Data represent mean values ± SEM. The gray dots indicate individual samples (N = 15): *, *P* < 0.05; ***, *P* < 0.001; Two-tailed unpaired *t*-tests were used, except that the result of RPS29 for *Cttnbp2^+/M120I^* mutant cultures was analyzed using a Mann-Whitney test. The light and dark blue dots indicate the averages of each batch of samples (N = 5); #, *P* < 0.05; ##, *P* < 0.01; ###, *P* < 0.001. Normalization indicates that the quantitative results of equal fractions were further normalized against the respective Coomassie Blue staining signals in Figures 7B and **7C**.

### Zinc treatment increases the neuronal population in the culture

Given that our culture conditions highly enrich neuronal populations (Liu *et al*., 2013), we expect the above-described observation of increased protein synthesis in our cultures to be primarily attributable to neurons. To confirm this point, we examined the cell population in our neuronal cultures by immunostaining for the neuronal marker MAP2 and the astrocyte/progenitor marker GFAP. The fluorescence images were randomly recorded blindly to quantify neuronal and non-neuronal populations (**Supplementary Figure S7**). We found that total cell numbers were slightly increased in the presence of an extra 100 μM zinc (**Supplementary** Figure 7**, S7C**). As expected, neurons proved to be the major cell type (80-85%) in our cultures, with GFAP-positive cells accounting for approximately 10-20% of both WT and *Cttnbp2^+/M120I^* cultures in the absence of extra zinc (**Supplementary Figure S7B, S7C**). Unexpectedly, in the presence of the extra 100 μM zinc, we found that the population of MAP2-positive cells was increased, accounting for over 90% of both WT and *Cttnbp2^+/M120I^* cultures, with GFAP-positive cells being reduced to less than 10% of both those culture types (**Supplementary Figure S7B, S7C**). A protein expression analysis by immunoblotting using antibodies recognizing SARM1 to represent a neuronal protein (Lin *et al*., 2014) and GFAP as a non-neuronal protein further indicated an increase in SARM1 proteins but a decrease in GFAP in cultures, regardless of genotype (**Supplementary Figure S8**). These data indicate that raising zinc levels in cultures increases the neuronal population, although we cannot be sure if this outcome is due to enhanced neurogenesis or neuronal survival. Importantly, the increment of protein synthesis in the presence of extra zinc is unlikely attributable to non-neuronal cells in the cultures.

### Long-term zinc supplementation restores the density and size of dendritic spines in male

#### Cttnbp2^+/M120I^ mice

Finally, we investigated if long-term dietary zinc supplementation endows any beneficial effects in terms of dendritic spine phenotypes in *Cttnbp2*-deficient mice. To do so, we provisioned mice with 30 or 150 ppm zinc diets starting from the embryonic stage to P42 (**Figure 9A**). To monitor dendritic spine morphology, we crossed *Cttnbp2^+/M120I^* mice to the Thy1-YFP-H line. Based on YFP signals, we measured the density and size of the dendritic spines of the infralimbic area of the medial prefrontal cortex (ILA), a brain region sensitive to *Cttnbp2* M120I mutation (Yen *et al*., 2023). Similar to our findings from a previous study on mice fed an 84-ppm zinc diet (Yen *et al*., 2023), we observed that *Cttnbp2^+/M120I^*mice fed on a 30-ppm zinc diet exhibited dendritic spine deficits, including reduced spine density, shorter spine length, and narrower spine width compared to wild-type (WT) littermates (**Figure 9B, 9C**). Importantly, *Cttnbp2^+/M120I^*mice fed on the 150-ppm zinc diet since the embryonic stage did not display a dendritic spine deficit, either in terms of density or size (**Figure 9B, 9C**), indicating that long-term zinc supplementation restores the dendritic spine deficits caused by *Cttnbp2* M120I mutation in male mice.

**Figure 9.**
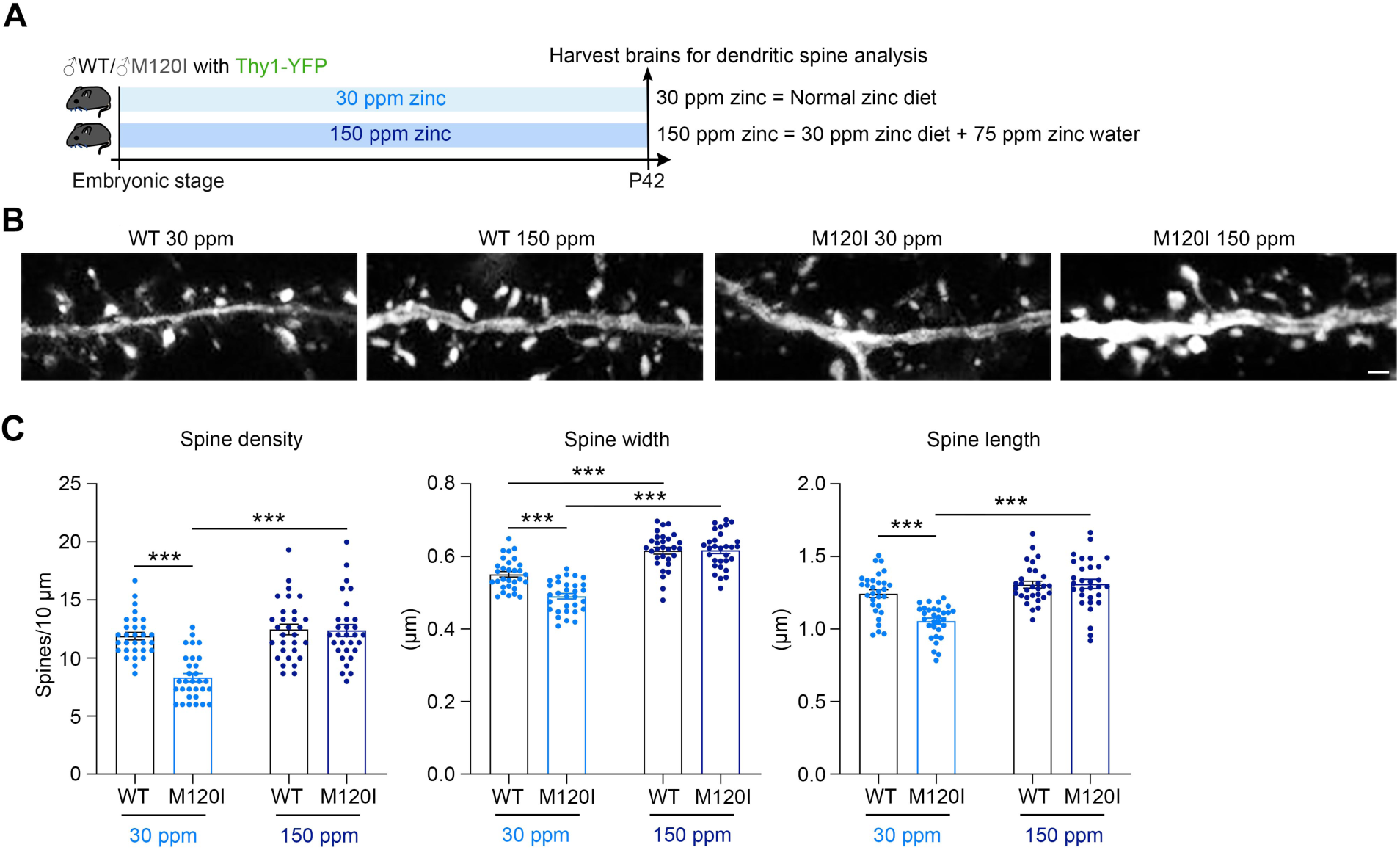
Long-term zinc supplementation rescues the synaptic morphology defects of male *Cttnbp2^+/M120I^*mice (**A**) Experimental condition for investigating the effect of long-term zinc supplementation on dendritic spine formation. Wild-type and *Cttnbp2^+/M120I^* mice were first crossed to Thy1-YFP mice. The YFP signals were used to outline the dendrite morphology of projection neurons at P42. Mice were fed on diets with either 30 or 150 ppm zinc starting at the embryonic stage and throughout the experimental period. (**B**) Representative images of dendritic segments of ILA neurons. (**C**) Quantification of (**B**). Results for the density, length and width of dendritic spines are shown. A total of 30 neurons collected from three mouse brains were analyzed for each group. Data represent mean ± SEM. Individual data points are also shown. *** *P* < 0.001; two-way ANOVA with Bonferroni multiple comparison test. Scale bar: 1 μm.

## Discussion

In this study, we employed ASD-linked *Cttnbp2* mutant mice as a genetic model to explore sex-specific responses to dietary zinc supplementation and the underlying regulatory mechanisms. In male *Cttnbp2*^+/M120I^ mice, increased zinc intake enhanced ribosome biogenesis and protein synthesis, restoring dendritic spine density and size, subsequently improving social behaviors. In contrast, female mice demonstrate resilience in brain function, remaining unaffected by zinc intake levels or *Cttnbp2* genetic variation. This resilience may stem from higher mTOR activity and elevated levels of translation initiation factors in female brains, which may enhance protein synthesis and offer protection against zinc deprivation. Sustained zinc supplementation is critical, as social deficits re-emerged in male *Cttnbp2^+/M120I^* mice within one to two weeks of zinc withdrawal. However, female mice continued to resist the effects of both the *Cttnbp2* mutation and zinc deprivation, exhibiting no significant social deficits. These findings highlight the role of sex-differential physiological responses in shaping the interactions between dietary zinc and genetic variations associated with ASD, strengthening the role of nutrients in ASD (Hsueh, 2025).

Our bioinformatics analysis of existing data indicates that ∼18% of SFARI genes are either zinc-binding or zinc-related. For SFARI Score 1 genes, the percentage of zinc-related genes is further increased to 25%. Notably, ∼13% of SFARI genes (156 genes) encode zinc finger proteins, representing at least four-fold enrichment relative to their proportion (3%) in the human genome (Klug, 2010). Among these zinc finger proteins, significant GO terms include zinc finger transcriptional regulators, histone modifiers, and metabolic processes. These results indicate that, apart from proteins directly involved in neurotransmission and synaptic organization, zinc regulates transcription and metabolism to indirectly support synaptic responses.

This scenario aligns with prior research demonstrating roles for zinc in DNA replication, protein synthesis, and cell growth across tissues such as skeletal muscle, heart, bone, liver, and brain (Dørup and Clausen, 1991; Duerre *et al*., 1977; Ma and Yamaguchi, 2001; Nishi, 1996; Williams and Chesters, 1970; Yamaguchi *et al*., 1988). Our proteomic analysis reveals that long-term zinc supplementation promotes ribosome biogenesis and enhances protein synthesis in male *Cttnbp2^+/M120I^* mice. The synchronized expression of ribosomal proteins is likely mediated by zinc finger transcription factors, including SP1 and YY1, whose binding sites are conserved in the promoter regions of ribosomal protein genes (Perry, 2005; Petibon *et al*., 2021). This information further supports that the beneficial effects of long-term zinc supplementation are not limited to improving the function of CTTNBP2, but are also achieved by the broad spectrum activities of zinc in cells.

Approximately 30% of patients with ASD exhibit reduced zinc levels in their bodies (Alsufiani *et al*., 2022; Arora *et al*., 2017; Yasuda *et al*., 2011). Given zinc’s essential role in neuronal functions, zinc supplementation may offer a potential treatment for alleviating ASD symptoms. In addition, a decrease of ∼70% in dendritic spine density, a common feature of ASD, has been estimated based on findings from ASD mouse models (Lu and Hsueh, 2022). Zinc supplementation may help counteract this deficit. Indeed, apart from the *Cttnbp2* mutation analyzed in this study, zinc supplementation has demonstrated beneficial effects in other ASD mouse models, including *Tbr1^+/–^* mice (Lee *et al*., 2022a), *Shank3* mutant mice (Fourie *et al*., 2018; Vyas *et al*., 2020), and BTBR mice (Zhang *et al*., 2023). These results underscore the potential of zinc as a broadly applicable therapeutic strategy for ASD (Hsueh, 2025).

It remains unclear why male mutant mice are particularly susceptible to zinc deprivation. Males generally require more zinc to maintain their health (Haase *et al*., 2020), which may intensify the competition for zinc between the brain and peripheral tissues. Since CTTNBP2 is specifically expressed in the brain, its ability to bind zinc plays a role in maintaining adequate zinc levels within the brain (Shih *et al*., 2020a). A deficiency in CTTNBP2 could disrupt zinc homeostasis in the brain and exacerbate overall zinc demand in affected mice. In this scenario, two potential strategies could mitigate the observed brain phenotypes: reducing peripheral zinc demand or increasing total zinc intake. The former is challenging to investigate experimentally. In this study, we have demonstrated that increasing zinc intake effectively improves pathological phenotypes of ASD.

Previous studies have indicated that protein synthesis by males and females are somewhat differentially regulated. For instance, estrogen activates protein synthesis in mouse uterine epithelial cells via the ERK-mTOR pathway (Wang *et al*., 2015). Moreover, estrogen receptor signaling and metabotropic glutamate receptor 5 (mGluR5) work together to enhance protein synthesis in female *Pten* knockout neurons (Molinaro *et al*., 2024). Our previous study also indicated that mTOR pathway activity is higher in female *Cttnbp2^+/M120I^*mice compared to male *Cttnbp2^+/M120I^* mice (Yen *et al*., 2023), potentially leading to higher protein synthesis and protective effects in female mice. This outcome also indicates that male *Cttnbp2^+/M120I^* mice may not effectively promote protein synthesis. Based on this notion, in a previous study, we stimulated the mTOR pathway in male *Cttnbp2^+/M120I^* mice by providing them with more branched-chain amino acids (BCAA) to investigate if protein synthesis is relevant to the deficits caused by *Cttnbp2* mutations (Yen *et al*., 2023). Notably, we found that BCAA supplementation for 1 week improved neuronal activation upon social stimulation, the dendritic spine density of neurons, and the social behaviors of male *Cttnbp2^+/M120I^* mice (Yen *et al*., 2023). In addition to *Cttnbp2* mutant mice, our previous studies have shown that BCAA supplementation also benefits *Vcp* mutant mice and *Nf1^+/–^*mice (Huang *et al*., 2021; Shih and Hsueh, 2016; Shih *et al*., 2020c). In the current study, we have shown that long-term zinc supplementation promotes ribosome biogenesis and protein synthesis. Thus, there is considerable crosstalk between how these two nutrient pathways, i.e., BCAA and zinc, promote protein synthesis to control dendritic spine formation and brain function. It would be intriguing to study the effect of zinc and BCAA on other ASD models and even patients.

It is well known that the gut-brain axis is relevant to ASD (Fattorusso *et al*., 2019; Fung *et al*., 2017; Kim *et al*., 2017; Korteniemi *et al*., 2023; Morton *et al*., 2023; Sharon *et al*., 2019; Willyard, 2021; Yap *et al*., 2021). Gut microbiota may be changed by diet. A study using 3D intestinal organoids and a mouse model have shown that zinc alters gut development and the gut microbiome (Sauer *et al*., 2021). Though our study using cultured neurons demonstrates that zinc acts directly on neurons to control protein synthesis, it is possible that long-term zinc supplementation also influences gut microbiota and contributes to the regulation of brain function, which warrants further study in the future.

## Supplementary information

Supplementary information on this work is available online.

## Acknowledgments

We thank the Bioinformatics Core and the Animal Facility of the Institute of Molecular Biology, Academia Sinica, for assistance, Dr. John O’Brien conducted English editing, and members of Y.-P.H.’s laboratory relabeled samples for blind experiments.

## Author contributions

Conceptualization, Y.-L.F., T.-L.Y., T.-F.W. and Y.-P.H.; Methodology and investigation, Y.-L.F., T.-L.Y., H.-C.L., T.-F.W. and Y.-P.H.; Writing, Y.-L.F., T.-L.Y., T.-F.W. and Y.-P.H.; Funding acquisition, Y.-P.H.; Supervision and project administration, Y.-P.H. All authors approved the submitted version.

## Materials Availability

This study did not generate unique or stable reagents.

## Data and Code Availability

This study did not generate any unique code. All data generated or analyzed during this study are included in this published article and its supplementary information files.

## References

Alsufiani, H.M., Alkhanbashi, A.S., Laswad, N.A.B., Bakhadher, K.K., Alghamdi, S.A., Tayeb, H.O., Tarazi, F.I., 2022. Zinc deficiency and supplementation in autism spectrum disorder and Phelan-McDermid syndrome. J Neurosci Res 100, 970–978.

Arora, M., Reichenberg, A., Willfors, C., Austin, C., Gennings, C., Berggren, S., Lichtenstein, P., Anckarsäter, H., Tammimies, K., Bölte, S., 2017. Fetal and postnatal metal dysregulation in autism. Nature Communications 8, 15493.

Baron, M.K., Boeckers, T.M., Vaida, B., Faham, S., Gingery, M., Sawaya, M.R., Salyer, D., Gundelfinger, E.D., Bowie, J.U., 2006. An architectural framework that may lie at the core of the postsynaptic density. Science 311, 531–535.

Brown, K.H., Peerson, J.M., Rivera, J., Allen, L.H., 2002. Effect of supplemental zinc on the growth and serum zinc concentrations of prepubertal children: a meta-analysis of randomized controlled trials. Am J Clin Nutr 75, 1062–1071.

Cassandri, M., Smirnov, A., Novelli, F., Pitolli, C., Agostini, M., Malewicz, M., Melino, G., Raschellà, G., 2017. Zinc-finger proteins in health and disease. Cell Death Discovery 3, 17071.

Chen, Y.K., Hsueh, Y.P., 2012. Cortactin-binding protein 2 modulates the mobility of cortactin and regulates dendritic spine formation and maintenance. J Neurosci 32, 1043–1055.

Cole, T.B., Wenzel, H.J., Kafer, K.E., Schwartzkroin, P.A., Palmiter, R.D., 1999. Elimination of zinc from synaptic vesicles in the intact mouse brain by disruption of the ZnT 3 gene. Proceedings of the national academy of sciences 96, 1716–1721.

De Rubeis, S., He, X., Goldberg, A.P., Poultney, C.S., Samocha, K., Cicek, A.E., Kou, Y., Liu, L., Fromer, M., Walker, S., Singh, T., Klei, L., Kosmicki, J., Shih-Chen, F., Aleksic, B., Biscaldi, M., Bolton, P.F., Brownfeld, J.M., Cai, J., Campbell, N.G., Carracedo, A., Chahrour, M.H., Chiocchetti, A.G., Coon, H., Crawford, E.L., Curran, S.R., Dawson, G., Duketis, E., Fernandez, B.A., Gallagher, L., Geller, E., Guter, S.J., Hill, R.S., Ionita-Laza, J., Jimenz Gonzalez, P., Kilpinen, H., Klauck, S.M., Kolevzon, A., Lee, I., Lei, I., Lei, J., Lehtimaki, T., Lin, C.F., Ma’ayan, A., Marshall, C.R., McInnes, A.L., Neale, B., Owen, M.J., Ozaki, N., Parellada, M., Parr, J.R., Purcell, S., Puura, K., Rajagopalan, D., Rehnstrom, K., Reichenberg, A., Sabo, A., Sachse, M., Sanders, S.J., Schafer, C., Schulte-Ruther, M., Skuse, D., Stevens, C., Szatmari, P., Tammimies, K., Valladares, O., Voran, A., Li-San, W., Weiss, L.A., Willsey, A.J., Yu, T.W., Yuen, R.K., Cook, E.H., Freitag, C.M., Gill, M., Hultman, C.M., Lehner, T., Palotie, A., Schellenberg, G.D., Sklar, P., State, M.W., Sutcliffe, J.S., Walsh, C.A., Scherer, S.W., Zwick, M.E., Barett, J.C., Cutler, D.J., Roeder, K., Devlin, B., Daly, M.J., Buxbaum, J.D., 2014. Synaptic, transcriptional and chromatin genes disrupted in autism. Nature 515, 209–215.

Dørup, I., Clausen, T., 1991. Effects of magnesium and zinc deficiencies on growth and protein synthesis in skeletal muscle and the heart. Br J Nutr 66, 493–504.

Duerre, J.A., Ford, K.M., Sandstead, H.H., 1977. Effect of zinc deficiency on protein synthesis in brain and liver of suckling rats. J Nutr 107, 1082–1093.

Fattorusso, A., Di Genova, L., Dell’Isola, G.B., Mencaroni, E., Esposito, S., 2019. Autism Spectrum Disorders and the Gut Microbiota. Nutrients 11.

Figiel, M., Górka, A.K., Górecki, A., 2023a. Zinc Ions Modulate YY1 Activity: Relevance in Carcinogenesis. Cancers (Basel) 15.

Figiel, M., Szubert, F., Luchinat, E., Bonarek, P., Baranowska, A., Wajda-Nikiel, K., Wilamowski, M., Miłek, P., Dziedzicka-Wasylewska, M., Banci, L., Górecki, A., 2023b. Zinc controls operator affinity of human transcription factor YY1 by mediating dimerization via its N-terminal region. Biochimica et Biophysica Acta (BBA) - Gene Regulatory Mechanisms 1866, 194905.

Fourie, C., Vyas, Y., Lee, K., Jung, Y., Garner, C.C., Montgomery, J.M., 2018. Dietary Zinc Supplementation Prevents Autism Related Behaviors and Striatal Synaptic Dysfunction in Shank3 Exon 13-16 Mutant Mice. Front Cell Neurosci 12, 374.

Frederickson, C.J. 1989. Neurobiology of Zinc and Zinc-Containing Neurons. In: International Review of Neurobiology. pp. 145-238. Eds. J.R. Smythies, R.J. Bradley. Academic Press.

Frederickson, C.J., Suh, S.W., Silva, D., Frederickson, C.J., Thompson, R.B., 2000. Importance of zinc in the central nervous system: the zinc-containing neuron. J Nutr 130, 1471s–1483s.

Fung, T.C., Olson, C.A., Hsiao, E.Y., 2017. Interactions between the microbiota, immune and nervous systems in health and disease. Nat Neurosci 20, 145–155.

Grabrucker, A.M., 2014. A role for synaptic zinc in ProSAP/Shank PSD scaffold malformation in autism spectrum disorders. Dev Neurobiol 74, 136–146.

Grabrucker, S., Jannetti, L., Eckert, M., Gaub, S., Chhabra, R., Pfaender, S., Mangus, K., Reddy, P.P., Rankovic, V., Schmeisser, M.J., Kreutz, M.R., Ehret, G., Boeckers, T.M., Grabrucker, A.M., 2014. Zinc deficiency dysregulates the synaptic ProSAP/Shank scaffold and might contribute to autism spectrum disorders. Brain 137, 137–152.

Haase, H., Ellinger, S., Linseisen, J., Neuhäuser-Berthold, M., Richter, M., 2020. Revised D-A-CH-reference values for the intake of zinc. Journal of Trace Elements in Medicine and Biology 61, 126536.

Hsueh, Y.P., 2025. Signaling in autism: Relevance to nutrients and sex. Curr Opin Neurobiol 90, 102962.

Hu, H.T., Shih, P.Y., Shih, Y.T., Hsueh, Y.P., 2016. The Involvement of Neuron-Specific Factors in Dendritic Spinogenesis: Molecular Regulation and Association with Neurological Disorders. Neural Plast 2016, 5136286.

Huang, E.P., 1997. Metal ions and synaptic transmission: think zinc. Proceedings of the National Academy of Sciences 94, 13386–13387.

Huang, T.N., Chuang, H.C., Chou, W.H., Chen, C.Y., Wang, H.F., Chou, S.J., Hsueh, Y.P., 2014. Tbr1 haploinsufficiency impairs amygdalar axonal projections and results in cognitive abnormality. Nat Neurosci 17, 240–247.

Huang, T.N., Shih, Y.T., Lin, S.C., Hsueh, Y.P., 2021. Social behaviors and contextual memory of Vcp mutant mice are sensitive to nutrition and can be ameliorated by amino acid supplementation. iScience 24, 101949.

Hung, Y.F., Chen, C.Y., Shih, Y.C., Liu, H.Y., Huang, C.M., Hsueh, Y.P., 2018. Endosomal TLR3, TLR7, and TLR8 control neuronal morphology through different transcriptional programs. J Cell Biol 217, 2727–2742.

Kim, S., Kim, H., Yim, Y.S., Ha, S., Atarashi, K., Tan, T.G., Longman, R.S., Honda, K., Littman, D.R., Choi, G.B., Huh, J.R., 2017. Maternal gut bacteria promote neurodevelopmental abnormalities in mouse offspring. Nature 549, 528–532.

Klug, A., 2010. The discovery of zinc fingers and their applications in gene regulation and genome manipulation. Annu Rev Biochem 79, 213–231.

Klug, A., Rhodes, D., 1987. Zinc fingers: a novel protein fold for nucleic acid recognition. Cold Spring Harb Symp Quant Biol 52, 473–482.

Korteniemi, J., Karlsson, L., Aatsinki, A., 2023. Systematic review: Autism spectrum disorder and the gut microbiota. Acta Psychiatr Scand 148, 242–254.

Lee, E.J., Lee, H., Huang, T.N., Chung, C., Shin, W., Kim, K., Koh, J.Y., Hsueh, Y.P., Kim, E., 2015. Trans-synaptic zinc mobilization improves social interaction in two mouse models of autism through NMDAR activation. Nat Commun 6, 7168.

Lee, K., Jung, Y., Vyas, Y., Skelton, I., Abraham, W.C., Hsueh, Y.P., Montgomery, J.M., 2022a. Dietary zinc supplementation rescues fear-based learning and synaptic function in the Tbr1(+/-) mouse model of autism spectrum disorders. Mol Autism 13, 13.

Lee, K., Mills, Z., Cheung, P., Cheyne, J.E., Montgomery, J.M., 2022b. The Role of Zinc and NMDA Receptors in Autism Spectrum Disorders. Pharmaceuticals (Basel) 16.

Lin, C.W., Liu, H.Y., Chen, C.Y., Hsueh, Y.P., 2014. Neuronally-expressed Sarm1 regulates expression of inflammatory and antiviral cytokines in brains. Innate Immun 20, 161–172.

Lin, C.W., Septyaningtrias, D.E., Chao, H.W., Konda, M., Atarashi, K., Takeshita, K., Tamada, K., Nomura, J., Sasagawa, Y., Tanaka, K., Nikaido, I., Honda, K., McHugh, T.J., Takumi, T., 2022. A common epigenetic mechanism across different cellular origins underlies systemic immune dysregulation in an idiopathic autism mouse model. Mol Psychiatry 27, 3343–3354.

Lin, Y.J., Huang, T.N., Hsueh, Y.P., 2023. Quantification of the density and morphology of dendritic spines and synaptic protein distribution using Thy1-YFP transgenic mice. STAR Protoc 4, 102290.

Liou, S.T., Wang, C., 2005. Small glutamine-rich tetratricopeptide repeat-containing protein is composed of three structural units with distinct functions. Arch Biochem Biophys 435, 253–263.

Liu, H.Y., Hong, Y.F., Huang, C.M., Chen, C.Y., Huang, T.N., Hsueh, Y.P., 2013. TLR7 negatively regulates dendrite outgrowth through the Myd88-c-Fos-IL-6 pathway. J Neurosci 33, 11479–11493.

Lu, M.H., Hsueh, Y.P., 2022. Protein synthesis as a modifiable target for autism-related dendritic spine pathophysiologies. Febs j 289, 2282–2300.

Ma, Z.J., Yamaguchi, M., 2001. Role of endogenous zinc in the enhancement of bone protein synthesis associated with bone growth of newborn rats. J Bone Miner Metab 19, 38–44.

Maenner, M., Warren, Z., Williams, A., al., e., 2023. Prevalence and Characteristics of Autism Spectrum Disorder Among Children Aged 8 Years — Autism and Developmental Disabilities Monitoring Network, 11 Sites, United States, 2020. MMWR Surveillance Summaries 72, 1–14.

Mocchegiani, E., Bertoni-Freddari, C., Marcellini, F., Malavolta, M., 2005. Brain, aging and neurodegeneration: role of zinc ion availability. Prog Neurobiol 75, 367–390.

Mohebalizadeh, M., Babapour, G., Maleki Aghdam, M., Mohammadi, T., Jafari, R., Shafiei-Irannejad, V., 2023. Role of Maternal Immune Factors in Neuroimmunology of Brain Development. Molecular Neurobiology.

Molinaro, G., Bowles, J.E., Croom, K., Gonzalez, D., Mirjafary, S., Birnbaum, S.G., Razak, K.A., Gibson, J.R., Huber, K.M., 2024. Female-specific dysfunction of sensory neocortical circuits in a mouse model of autism mediated by mGluR5 and estrogen receptor α. Cell Rep 43, 114056.

Mony, L., Paoletti, P., 2023. Mechanisms of NMDA receptor regulation. Curr Opin Neurobiol 83, 102815.

Morton, J.T., Jin, D.-M., Mills, R.H., Shao, Y., Rahman, G., McDonald, D., Zhu, Q., Balaban, M., Jiang, Y., Cantrell, K., Gonzalez, A., Carmel, J., Frankiensztajn, L.M., Martin-Brevet, S., Berding, K., Needham, B.D., Zurita, M.F., David, M., Averina, O.V., Kovtun, A.S., Noto, A., Mussap, M., Wang, M., Frank, D.N., Li, E., Zhou, W., Fanos, V., Danilenko, V.N., Wall, D.P., Cárdenas, P., Baldeón, M.E., Jacquemont, S., Koren, O., Elliott, E., Xavier, R.J., Mazmanian, S.K., Knight, R., Gilbert, J.A., Donovan, S.M., Lawley, T.D., Carpenter,, Bonneau, R., Taroncher-Oldenburg, G., 2023. Multi-level analysis of the gut–brain axis shows autism spectrum disorder-associated molecular and microbial profiles. Nature Neuroscience 26, 1208–1217.

Nishi, Y., 1996. Zinc and growth. J Am Coll Nutr 15, 340–344.

Ohoka, Y., Takai, Y., 1998. Isolation and characterization of cortactin isoforms and a novel cortactin-binding protein, CBP90. Genes Cells 3, 603-612.

Perry, R.P., 2005. The architecture of mammalian ribosomal protein promoters. BMC Evolutionary Biology 5, 15.

Petibon, C., Malik Ghulam, M., Catala, M., Abou Elela, S., 2021. Regulation of ribosomal protein genes: An ordered anarchy. Wiley Interdiscip Rev RNA 12, e1632.

Pfaender, S., Sauer, A.K., Hagmeyer, S., Mangus, K., Linta, L., Liebau, S., Bockmann, J., Huguet, G., Bourgeron, T., Boeckers, T.M., Grabrucker, A.M., 2017. Zinc deficiency and low enterocyte zinc transporter expression in human patients with autism related mutations in SHANK3. Sci Rep 7, 45190.

Portbury, S.D., Adlard, P.A., 2017. Zinc Signal in Brain Diseases. Int J Mol Sci 18, 2506.

Sanders, S.J., He, X., Willsey, A.J., Ercan-Sencicek, A.G., Samocha, K.E., Cicek, A.E., Murtha, M.T., Bal, V.H., Bishop, S.L., Dong, S., Goldberg, A.P., Jinlu, C., Keaney, J.F., 3rd, Klei, L., Mandell, J.D., Moreno-De-Luca, D., Poultney, C.S., Robinson, E.B., Smith, L., Solli-Nowlan, T., Su, M.Y., Teran, N.A., Walker, M.F., Werling, D.M., Beaudet, A.L., Cantor, R.M., Fombonne, E., Geschwind, D.H., Grice, D.E., Lord, C., Lowe, J.K., Mane, S.M., Martin, D.M., Morrow, E.M., Talkowski, M.E., Sutcliffe, J.S., Walsh, C.A., Yu, T.W., Ledbetter, D.H., Martin, C.L., Cook, E.H., Buxbaum, J.D., Daly, M.J., Devlin, B., Roeder, K., State, M.W., 2015. Insights into Autism Spectrum Disorder Genomic Architecture and Biology from 71 Risk Loci. Neuron 87, 1215–1233.

Sauer, A.K., Malijauskaite, S., Meleady, P., Boeckers, T.M., McGourty, K., Grabrucker, A.M., 2021. Zinc is a key regulator of gastrointestinal development, microbiota composition and inflammation with relevance for autism spectrum disorders. Cell Mol Life Sci 79, 46.

Sharon, G., Cruz, N.J., Kang, D.W., Gandal, M.J., Wang, B., Kim, Y.M., Zink, E.M., Casey, C.P., Taylor, B.C., Lane, C.J., Bramer, L.M., Isern, N.G., Hoyt, D.W., Noecker, C., Sweredoski, M.J., Moradian, A., Borenstein, E., Jansson, J.K., Knight, R., Metz, T.O., Lois, C., Geschwind, D.H., Krajmalnik-Brown, R., Mazmanian, S.K., 2019. Human Gut Microbiota from Autism Spectrum Disorder Promote Behavioral Symptoms in Mice. Cell 177, 1600–1618.e1617.

Shevchenko, A., Tomas, H., Havlis, J., Olsen, J.V., Mann, M., 2006. In-gel digestion for mass spectrometric characterization of proteins and proteomes. Nat Protoc 1, 2856–2860.

Shih, P.Y., Fang, Y.L., Shankar, S., Lee, S.P., Hu, H.T., Chen, H., Wang, T.F., Hsia, K.C., Hsueh, Y.P., 2022. Phase separation and zinc-induced transition modulate synaptic distribution and association of autism-linked CTTNBP2 and SHANK3. Nat Commun 13, 2664.

Shih, P.Y., Hsieh, B.Y., Lin, M.H., Huang, T.N., Tsai, C.Y., Pong, W.L., Lee, S.P., Hsueh, Y.P., 2020a. CTTNBP2 Controls Synaptic Expression of Zinc-Related Autism-Associated Proteins and Regulates Synapse Formation and Autism-like Behaviors. Cell Rep 31, 107700.

Shih, P.Y., Hsieh, B.Y., Tsai, C.Y., Lo, C.A., Chen, B.E., Hsueh, Y.P., 2020b. Autism-linked mutations of CTTNBP2 reduce social interaction and impair dendritic spine formation via diverse mechanisms. Acta Neuropathol Commun 8, 185.

Shih, Y.T., Hsueh, Y.P., 2016. VCP and ATL1 regulate endoplasmic reticulum and protein synthesis for dendritic spine formation. Nat Commun 7, 11020.

Shih, Y.T., Huang, T.N., Hu, H.T., Yen, T.L., Hsueh, Y.P., 2020c. Vcp Overexpression and Leucine Supplementation Increase Protein Synthesis and Improve Fear Memory and Social Interaction of Nf1 Mutant Mice. Cell Rep 31, 107835.

Takeda, A., 2001. Zinc homeostasis and functions of zinc in the brain. Biometals 14, 343–351.

Torigoe, T., Izumi, H., Yoshida, Y., Ishiguchi, H., Okamoto, T., Itoh, H., Kohno, K., 2003. Low pH enhances Sp1 DNA binding activity and interaction with TBP. Nucleic acids research 31, 4523–4530.

Tóth, K., 2011. Zinc in neurotransmission. Annual review of nutrition 31, 139–153.

Vorontsov, I.E., Fedorova, A.D., Yevshin, I.S., Sharipov, R.N., Kolpakov, F.A., Makeev, V.J., Kulakovskiy, I.V., 2018. Genome-wide map of human and mouse transcription factor binding sites aggregated from ChIP-Seq data. BMC Res Notes 11, 756.

Vyas, Y., Jung, Y., Lee, K., Garner, C.C., Montgomery, J.M., 2021. In vitro zinc supplementation alters synaptic deficits caused by autism spectrum disorder-associated Shank2 point mutations in hippocampal neurons. Mol Brain 14, 95.

Vyas, Y., Lee, K., Jung, Y., Montgomery, J.M., 2020. Influence of maternal zinc supplementation on the development of autism-associated behavioural and synaptic deficits in offspring Shank3-knockout mice. Mol Brain 13, 110.

Wang, Y., Zhu, L., Kuokkanen, S., Pollard, J.W., 2015. Activation of protein synthesis in mouse uterine epithelial cells by estradiol-17β is mediated by a PKC-ERK1/2-mTOR signaling pathway. Proc Natl Acad Sci U S A 112, E1382–1391.

Williams, R.B., Chesters, J.K., 1970. The effects of early zinc deficiency on DNA and protein synthesis in the rat. Br J Nutr 24, 1053–1059.

Willyard, C., 2021. How gut microbes could drive brain disorders. Nature 590, 22–25.

Yamaguchi, M., Oishi, H., Suketa, Y., 1988. Zinc stimulation of bone protein synthesis in tissue culture. Activation of aminoacyl-tRNA synthetase. Biochem Pharmacol 37, 4075–4080.

Yap, C.X., Henders, A.K., Alvares, G.A., Wood, D.L.A., Krause, L., Tyson, G.W., Restuadi, R., Wallace, L., McLaren, T., Hansell, N.K., Cleary, D., Grove, R., Hafekost, C., Harun, A., Holdsworth, H., Jellett, R., Khan, F., Lawson, L.P., Leslie, J., Frenk, M.L., Masi, A., Mathew, N.E., Muniandy, M., Nothard, M., Miller, J.L., Nunn, L., Holtmann, G., Strike, L.T., de Zubicaray, G.I., Thompson, P.M., McMahon, K.L., Wright, M.J., Visscher, P.M., Dawson, P.A., Dissanayake, C., Eapen, V., Heussler, H.S., McRae, A.F., Whitehouse, A.J.O., Wray, N.R., Gratten, J., 2021. Autism-related dietary preferences mediate autism-gut microbiome associations. Cell 184, 5916–5931.e5917.

Yasuda, H., Yoshida, K., Yasuda, Y., Tsutsui, T., 2011. Infantile zinc deficiency: association with autism spectrum disorders. Sci Rep 1, 129.

Yen, T.L., Huang, T.N., Lin, M.H., Hsu, T.T., Lu, M.H., Shih, P.Y., Ellegood, J., Lerch, J., Hsueh, Y.P., 2023. Sex bias in social deficits, neural circuits and nutrient demand in Cttnbp2 autism models. Brain 146, 2612–2626.

Zhang, L., Xu, X., Ma, L., Wang, X., Jin, M., Li, L., Ni, H., 2023. Zinc Water Prevents Autism-Like Behaviors in the BTBR Mice. Biol Trace Elem Res 201, 4779–4792.

